# Real-time measurement of short-chain fatty acids via microwave sensing: A pilot study

**DOI:** 10.64898/2026.02.10.705113

**Authors:** Kelsey Pinera, Judy Malas, Sebastian Reczek, Kiersten Oderman, Jamie Kaiser, Joe Walker, Dieff Vital, Jarrad Hampton-Marcell

**Author notes:** Corresponding author Jarrad Hampton-Marcell, Assistant Professor, Department of Biological Sciences, University of Illinois Chicago, 950 S. Halsted St. Chicago, IL 60607. denotes co-first authors (Kelsey Pinera, Judy Malas).

## Abstract

**Background:** Short-chain fatty acids (SCFAs) - acetate, propionate, and butyrate - are key microbial metabolites that influence host metabolism, barrier function, and immune tone. Yet, in vivo measurements of SCFAs remain poorly characterized because stool measurements only capture a small, spatially averaged fraction of luminal production.

**Purpose:** To evaluate whether label-free microwave sensing using a vector network analyzer (VNA) can differentiate SCFA identity, concentration, and their ratios via unique dielectric signatures.

**Methods:** Sodium acetate, sodium propionate, and sodium butyrate solutions (10 uM - 1M) were prepared by measuring their electrochemical properties (pH and mV). Using a 2-port VNA interfaced with a high-frequency copper plate (1 - 3 GHz), scattering and impedance parameters were measured for SCFAs and their ratios. A per-frequency ANOVA was employed to identify discriminatory frequency bands, and supervised machine learning (random forest) was employed to identify additional features differentiating SCFAs.

**Results:** SCFAs displayed distinct (ANOVA, p < 0.05) electrochemical signatures for both pH and mV. The 21 transmission spectrum revealed discriminatory frequencies with prominent bands at 1.38 - 1.40 GHz and 1.79 - 1.85 GHz. For SCFA ratios, the random forest model achieved up to 80.6% accuracy (k = 0.72) with S11 (77.7%) and S22 (69.9%) unwrapped phases observed as the top two important features contributing to the model’s performance.

**Conclusion:** VNA-based microwave sensing resolves SCFA-specific, frequency-dependent dielectric signatures across biologically relevant concentrations and classifies SCFA-like mixtures with high accuracy. These findings support microwave sensing as a foundation for real-time, non-invasive monitoring of gut microbial metabolism.

**IMPORTANCE:** Short-chain fatty acids (SCFAs) are central to host-microbe interactions, yet conventional stool assays provide only snapshots of microbial metabolism and poorly reflect real-time intestinal production of these microbial metabolites. SCFAs mediate intestinal barrier integrity, immune regulation, energy availability, and energy metabolism. Alterations in SCFA production and composition are increasingly recognized as biomarkers of gut microbial dysbiosis, with strong connections to conditions such as inflammatory bowel disease, metabolic syndrome, obesity, and neurodegenerative disorders. As such, the dynamic and quantitative assessment of SCFA profiles in vivo may provide valuable insight into gut health, disease progression, and therapeutic response. Despite large technological advances in the past decade, there is a limited understanding of in vivo SCFA dynamics within the gut, largely due to the inaccessibility and limitation of developing non-invasive diagnostic tools. Furthermore, the spatiotemporal resolution of microbial-derived metabolites in the gut is also very limited, with most current research methods relying on utilizing stool samples, which only contain a fraction of produced SCFA concentrations. In this pilot study, SCFA solutions of known concentrations and identity were measured and analyzed via their capacitance to use as an emerging diagnostic tool in clinical and/or athletic research and to further practice assessing gut microbial dynamics. We demonstrate that microwave sensing - a label-free, electrical approach - can distinguish among SCFAs and their concentrations by exploiting their dielectric differences and frequency-resolved behavior. Using a Vector Network Analyzer and a low-loss dielectric sensor, we identify reproducible GHz-scale bands that separate acetate, propionate, and butyrate. Results indicated that these metabolites can be differentiated via their S parameter (frequency response) electrical signals and pH values. Additionally, we classify physiological SCFA mixtures with strong performance.The study will be repeated by measuring various fluid media, starting with acidic solutions and testing with fluid mixes such as mock carbohydrate solutions that best mimic gut microbial conditions in hopes of developing a diagnostic tool (i.e., swallowable sensor) that can be used to monitor real-time human gut environments. This establishes a practical path toward dynamic, non-invasive readouts of gut microbial activity with potential applications in clinical monitoring.

## INTRODUCTION

Cardiometabolic disease (CMD) is the leading cause of death globally, accounting for approximately 17.9 million deaths annually (WHO, 2019). In recent years, a growing body of evidence has linked the microorganisms inhabiting the gastrointestinal (GI) tract (i.e., the gut microbiome) to host metabolic and inflammatory pathways relevant to CMD risk (Nesci et al., 2023). Among microbiome-derived mediators, short chain fatty acids (SCFAs) - primarily acetate, propionate, and butyrate - have been been linked to host health acting as both energy substrates and signaling molecules implicated in diabetes, obesity, hypertension and other CMDs (den Besten et al., 2013). Specifically, SCFAs are capable of influencing intestinal barrier integrity, blood pressure regulation, glucose and lipid homeostasis, and systemic inflammation implicating their importance and utility in shaping host-microbe interactions (Nogal et al., 2021). SCFAs are produced in different proportions depending on both substrate availability and the specific bacterial communities present in an individual’s gut microbiome. Different carbohydrate sources yield distinct SCFA profiles: for instance, resistant starch fermentation produces high levels of butyrate, while inulin fermentation generates substantial amounts of both butyrate and propionate (Baxter et al., 2019). Each SCFA exhibits unique physiological functions that contribute to host health. Acetate is often attributed to energy metabolism, propionate acts as a signaling molecule in the gut that supports metabolic and immune balance by stimulating gut hormones, maintaining the gut barrier, and reducing inflammation, and butyrate maintains the epithelial barrier and exerts anti-inflammatory effects that extend beyond the gut (Nogal et al., 2021)

Despite the recognized importance of SCFAs in human health, accurate measurement of these metabolites presents significant analytical challenges. Current methodologies for SCFA quantification rely primarily on ex vivo analysis of fecal samples using gas chromatography-mass spectrometry (GC-MS) or liquid chromatography-mass spectrometry (LC-MS). Mass spectrometry techniques enable precise identification and quantification of SCFAs by measuring their mass-to-charge ratios through sequential ionization, separation, and detection processes (Trivedi et al., 2022). While these analytical approaches have advanced our understanding of SCFA production and gut microbiome interactions, they possess inherent limitations. Fecal SCFA measurements represent only a small fraction (∼5%) of the total SCFAs produced in the intestinal lumen, as the majority are rapidly absorbed by the colonic epithelium (Primec et al., 2017). This means that relying solely on fecal analysis underestimates the true metabolic output of the gut microbiota. Furthermore, SCFA concentrations vary significantly along the GI tract, with higher concentrations typically observed in the proximal colon compared to distal regions, spatial information that is lost in fecal analysis (Primec et al., 2017). These limitations create challenges in accurately assessing the true physiological impact of SCFA production and their effects on host metabolism. As an alternative approach to attempt to measure SCFA concentrations that reflect physiological concentrations, in vitro gut microbiome models, including sophisticated bioreactor systems, have been created. These systems allow for a controlled manipulation of microbial communities and substrate availability (Zaarour et al., 2025), but they fail to capture key in vivo dynamics such as host-microbe interactions, immune responses, and physiological feedback mechanisms, which can significantly influence SCFA production and absorption.

These limitations have driven the development of non-invasive, real-time monitoring systems that provide readouts of intestinal biochemistry. Ingestible biosensors demonstrate feasibility for monitoring GI environments, and microwave sensing has emerged as a complementary, label-free approach that works by detecting changes in dielectric properties associated with solute composition without the use of any chemical tags and/or labels (Allaband et al., 2019). Biosensors capable of analyzing the GI environment and wirelessly communicating measurements to external devices have offered unprecedented insight into dynamic gut processes. Through implementation of microwave sensing via a vector network analyzer (VNA), it has been demonstrated that scattering parameters (S-parameters) can resolve frequency-dependent electro-magnetic behavior of solutions and tissues (Mimee et al., 2018). Notably, VNA has been employed to monitor metabolism by tracking the dielectric properties of blood glucose via changes in relative permittivity in human plasma (Juan et al., 2019). This new approach suggests a route to differentiate metabolites within biological mixtures by their dielectric properties. Furthermore, since SCFAs– primarily acetate, propionate, and butyrate– exhibit distinct dielectric potentials that arise from differences in molecular polarity, chain length, and respected pKa values, this allows them to produce a characteristic frequency-dependent electrical response.

In this study, we test whether microwave sensing can resolve SCFA identity and physiologically-inspired SCFA ratios. Using a VNA with a custom copper sensor, we (i) characterize electro-magnetic properties (pH, mV) of acetate, propionate, and butyrate, (ii) evaluate frequency-resolved S-parameters to discriminate among SCFAs, and (iii) apply machine learning to S-parameter features to classify mock carbohydrate conditions via their SCFA profiles. This study establishes a foundation for microwave sensing as a real-time, non-invasive window into gut microbial metabolism.

## METHODS

### Study overview

Solutions spanning 10 to 100uM (10uM increments) of acetate, propionate, or butyrate were generated. Following calibration using buffer solutions (4.01, 7.00, and 10.01), pH and millivolts (mV) were measured for each SCFA solution in triplicate. In addition to electro-chemical measurements, S-parameters were measured for SCFAs and mock carbohydrates using a vector network analyzer (LibreVNA) with a custom copper sensor plate and 3D-printed containment system.

### Reagents and SCFA solutions

1.0 M stock solutions of sodium acetate (MP Biomedical; 82.03 g/mol), sodium propionate (Thermo; 96.06 g/mol) and sodium butyrate (Thermo; 110.09 g/mol) were prepared by dissolving 1.231 g, 1.44 g, and 1.651 g, respectively, in molecular biology-grade water (Thermo). These solutions were serially diluted to generate 0.1 mM stock and working solutions of 10 - 100 uM. All solutions were vortexed after preparation and stored at room temperature until use. Mock carbohydrates including glucose, hydrolyzed guar gum, and hydrolyzed inulin solutions were generated mimicking the resultant ratios of acetate, propionate, and butyrate post-bacterial consumptions, according to den Besten et al. (2013).

### Electrochemical measurements

Before measurements, buffers at pH 4.01, 7.00, and 10.01 were vortexed and used to calibrate a glass pH electrode per manufacturer instructions (Fisher). SCFA solutions were equilibrated at room temperature, vortexed (1-20 s, ∼2500 rpm), and measured in triplicate. After each reading, the electrode was rinsed with distilled water and gently dried with lint-free wipes. For each sample, pH, mV, temperature, and meter status were recorded.

### Sensor plate

Sensor plates (Brigitflex, Elgin, IL) were generated using a high-frequency laminate (Rogers RO3010; dielectric constant Dk = 10.2, loss tangent tan δ = 0.0022, thickness = 1.28 mm) in order to provide a stable, low-loss dielectric for microwave interrogation in the 1-3 GHz band, while also minimizing spurious attenuation unrelated to analyte composition (**Supp. Figure 1**). Each copper plate was fitted with a 3D-printed PLA well (18.75 mm x 18.75 mm, 10 mm height) and bonded with gel cyanoacrylate, which included a 1.5mm brim to reduce leaks. To prevent cross-contamination and surface conditioning effects, a dedicated plate was used for each SCFA. Plates were rinsed thoroughly with molecular-grade water between measurements.

### Electromagnetic measurements

A 2-port VNA (LibreVNA) with frequency ranges of 100 kHz - 6 GHz was calibrated using a SOLT12 configuration according to the manufacturer’s suggested protocol. Measurements were acquired from 1 - 3 GHz to balance penetration depth and dielectric sensitivity while minimizing low-frequency ionic conduction artifacts. For each sample, 1mL was pipetted into the 3D-printed well; readings proceeded from lowest to highest concentrations with water flushes between samples. S-parameters and impedance-derived metrics were exported from an open-sourced LibreVNA-GUI software for further analysis. A total of 57 measures for each sample were recorded from the VNA (**Supplemental Table 1**). Among recorded measurements, key features - S11 magnitude, S11 unwrapped Phase, and S21 unwrapped phase - were configured and analyzed for further assessment. The S-parameters, S11 and S21 describe the reflection and transmission behavior of the microwave signal that interacts with the sample. S11 represents the fraction of the incident wave that is reflected back toward the source, while S21 represents the fraction that is transmitted through the sample. The magnitude of S11 represents the strength of the reflection and its unwrapped phase reveals changes in the dielectric response. Lastly, the unwrapped phase of S21 captures the phase delays (i.e. how long the frequency component of the sample takes to pass through the system) of the sample which reveals the sensitivity of the sample’s characteristics and dielectric constant. We posited that these three parameters together would enable the differentiation of SCFA metabolites based on their frequency-dependent electromagnetic responses.

### Statistical analysis

The R programming language was used with RStudio (version 4.5.0) for all statistical analyses. Frequency values were converted to gigahertz (GHz) for interpretability. To discern frequency bands that discriminated among SCFAs and mock carbohydrates, a per-frequency parametric analysis of variance (ANOVA) was conducted via the R package ‘rstatix’ (Kassambara, 2023). Frequencies (1.0 - 3.0 GHz), were divided into 50 MHz bins; bins with an adjusted p-value < 0.05 were considered significant following Benjamini-Hochberg correction to control for false-discovery rate (FDR). Significant frequency bands were visualized using-log_10_(p).

To identify informative electromagnetic features of mock carbohydrates, a random forest classifier was employed using the R package ‘caret’. Following filtering of zero-values and non-zero variance, the data was split into an 80% training dataset and 20% test dataset, ensuring electromagnetic features remained balanced. The model used 500 decision trees, a stratified 5-fold cross validation, and hyperparameter turning (e.g., 4, 8, 16, and 30 predictors for each split). The model’s performance was validated using overall accuracy, Cohen’s Kappa, and sensitivity and specificity via the confusion matrix. Important features were extracted from the random forest model using the R packages ‘caret’ and ‘randomForest’ to assess each variable’s contribution to the overall model’s accuracy. All visualizations were generated using the R packages ‘ggplot2’,’ggsci’,’ggpubr’, and ‘patchwork’ (Wickham, 2016; Xiao, 2025; Kassambara, 2025; Pedersen, 2025).

## RESULTS

### SCFAs are differentiated via their electrochemical and electromagnetic properties

To discern differences in electrochemical properties of SCFAs, pH and voltage (mV) were assessed for acetate, propionate, and butyrate solutions with concentrations between 10 uM - 100 uM. A Pearson correlation revealed significant (p < 0.05) inverse associations between concentration and pH for acetate (R = -0.66), butyrate (R = -0.77) and propionate ( R = -0.52) (**Supp. Figure 2A**). Wilcoxon rank test displayed that butyrate significantly differed from acetate in both pH (p = 0.02) and mV (p = 0.016) but there was no significant differences between acetate and propionate or butyrate and propionate.

To determine whether the S21 scattering parameter could distinguish between SCFAs, we analyzed the transmission coefficient across frequencies ranging from 0.9 to 3.0 GHz. A per-frequency one-way ANOVA was employed to identify specific frequencies at which the S21 signal significantly differed by SCFA type (**Supplemental Table 1**). Several frequency bands exhibited highly significant differences following false discovery rate correction (**Table 1**). The largest difference was observed in the S21 range between 1.79-1.85 GHz, peaking at 1.82 GHz (**Figure 2A**; ANOVA, p_FDR_ = 9.25 x 10^-14^). Additionally, statistically significant signals appeared between 1.38-1.40 GHz, suggesting multiple frequency distributions wherein SCFAs are differentially detected. This was further supported by overlaying the mean S21 traces for each SCFA with the standard error of the mean identifying significant frequencies (ANOVA, p < 0.05) (**Figure 2B**). Clear differences were observed for SCFAs, particularly between 1.3-1.4 GHz and 1.8-2.0 GHz. Importantly, this reveals a band-averaged distribution for the S21 spectrum within the 1.3-1.4 GHz range (**Figure 2C**) while the 1.8-2.0 GHz range (**Figure 2D**) had revealed significant (ANOVA, p < 0.001) differences. To add, butyrate produced greater signal attenuation (i.e., weakening or potential loss of the signal) relative to acetate and propionate within these frequencies. For example, at 1.8-2.0 GHz, butyrate had a significantly lower S21 parameter than acetate and propionate (p_FDR_ = 3 x 10^-14^, S21delta = -3.74 to -4.31 dB). This suggests that the dielectric properties of SCFAs result in distinct frequency-dependent electromagnetic behavioral shifts. Further, all SCFAs were significantly different at 1.8-2.0 GHz supporting the potential of a tripartite separation window (**Table 2**). Combined, these results suggest the S21 spectrum provides frequency-resolved sensitivity among SCFAs. Notably, the 1.8-2.0 GHz range is a potential window for differentiating the three major SCFAs.

**Figure 1.**
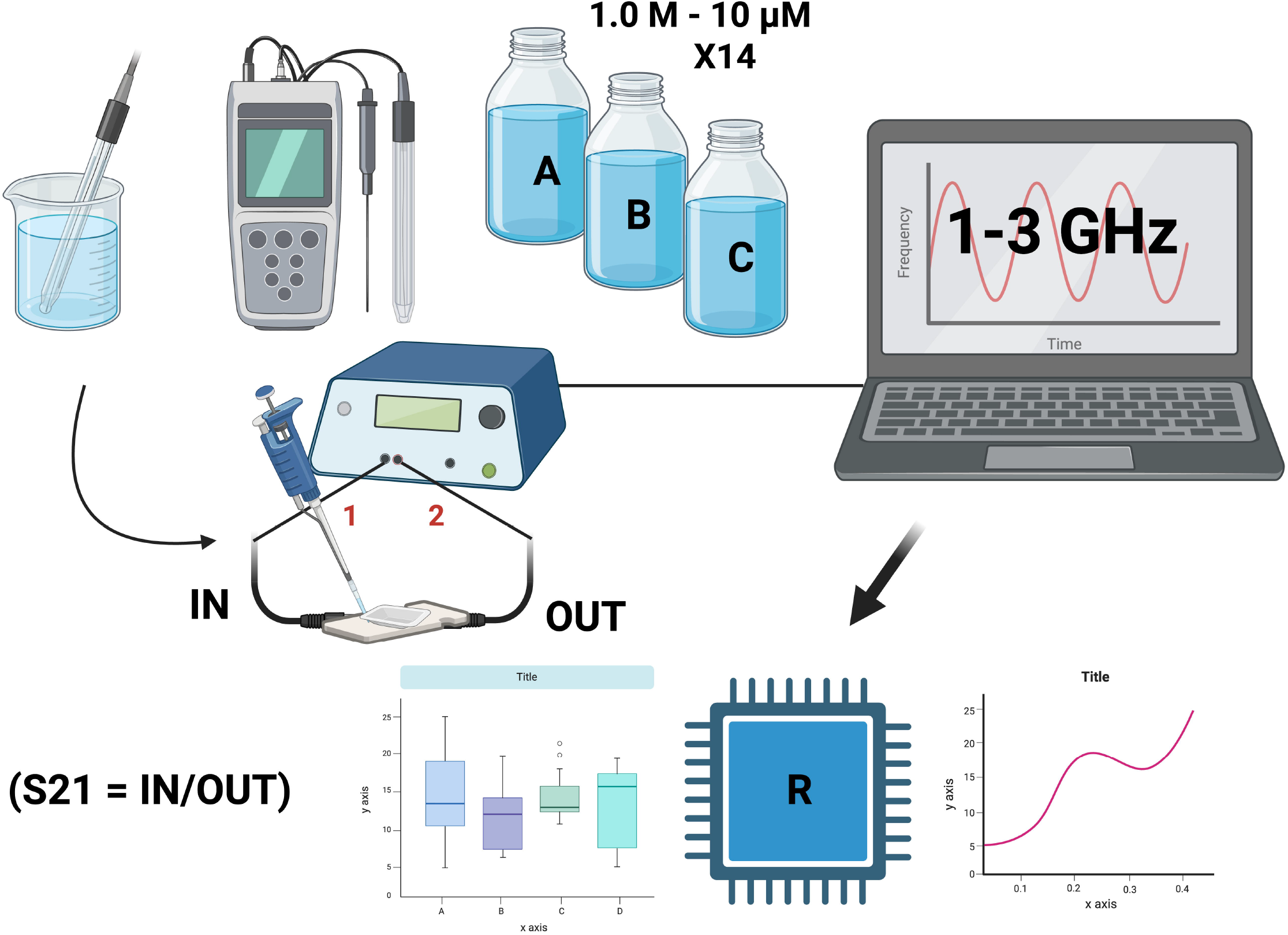
Experimental workflow. Acetate, propionate, and butyrate (10 uM - 1.0 M) were prepared and measured for pH/mV followed by measurements of their electromagnetic properties using a vector network analyzer. Scattering parameters (S-parameters) were analyzed in R to detect differences.

**Figure 2.**
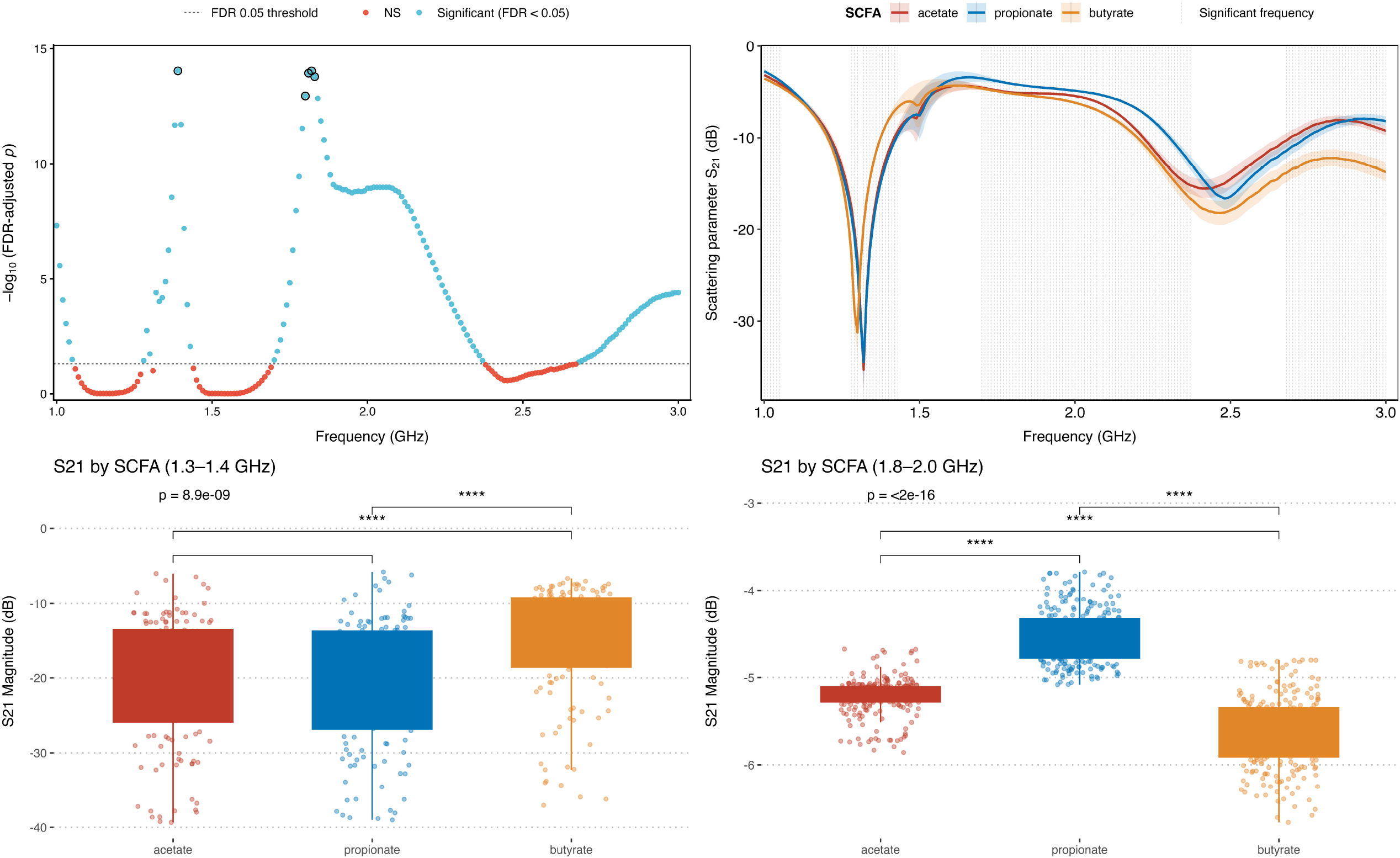
Frequency-resolved separation of SCFAs by transmission. (**A**) Per-frequency one-way ANOVA testing S21 differences among SCFAs. Points show-log10(p) following correction for false discovery rates; the dashed lines mark an alpha = 0.05. (**B**) Mean S21 traces by SCFA are depicted with vertical gray bands indicating significant (p < 0.05) regions. (**C-D**) Box and dot plots of S21 within two frequency bands (1.3-1.4 GHz and 1.8-2.0 GHz) are depicted. A pairwise Wilcoxon test with BH correction was performed. Butyrate shows greater attenuation than acetate/propionate within these bands.

**Table 1.** Frequency bands with significant S21 separation among SCFAs. A summary of per-frequency one-way ANOVAs across frequencies (1.0-3.0 GHz) binned at 50 MHz comparing S21-parameters among acetate, propionate, and butyrate.

| Frequency (GHz) | Adjusted P-value (ANOVA) | Number of Significant Comparisons | Significant Pairwise Comparisons |
| --- | --- | --- | --- |
| 1.39 | $9.25 \times 10^{-15}$ | 2 | acetate-butyrate;propionate-butyrate |
| 1.82 | $9.25 \times 10^{-15}$ | 3 | acetate-propionate; acetate-butyrate; propionate-butyrate |
| 1.81 | $9.25 \times 10^{-15}$ | 2 | acetate-propionate;propionate-butyrate |
| 1.83 | $9.25 \times 10^{-15}$ | 3 | acetate-propionate; acetate-butyrate; propionate-butyrate |
| 1.8 | $9.25 \times 10^{-15}$ | 2 | acetate-propionate; propionate-butyrate |
| 1.84 | $9.25 \times 10^{-15}$ | 3 | acetate-propionate; acetate-butyrate; propionate-butyrate |
| 1.85 | $9.25 \times 10^{-15}$ | 3 | acetate-propionate; acetate-butyrate; propionate-butyrate |
| 1.4 | $9.25 \times 10^{-15}$ | 2 | acetate-butyrate; propionate-butyrate |
| 1.38 | $9.25 \times 10^{-15}$ | 2 | acetate-butyrate; propionate-butyrate |
| 1.79 | $9.25 \times 10^{-15}$ | 2 | acetate-propionate; propionate-butyrate |

**Table 2.** SCFA production rates for mock carbohydrates. The values show the relative proportions of the major short-chain fatty acids for each of the four mock carbohydrates

| Substrate | SCFA Production Rate |  |  |
| --- | --- | --- | --- |
|  | Acetate | Propionate | Butyrate |
|  | mmol/L reaction volume/h |  |  |
| Glucose | 21.2 | 5.0 | 2.2 |
| Hydrolyzed Guar Gum | 13.3 | 11.1 | 1.6 |
| Hydrolyzed Inulin | 15.1 | 6.4 | 3.9 |
| Water (Control) | 0 | 0 | 0 |

### Machine learning reveal scattering parameters that distinguish SCFA ratios

To determine whether additional electromagnetic features outside of S21 could differentiate SC-FAs, supervised machine learning was employed on mock carbohydrate solutions with defined ratios of acetate, propionate, and butyrate that were distinctly different from one another (den Besten et al., 2013; **Table 2**). A random forest classifier was trained on 4,411 measurements each reporting 56 scattering parameters and impedance metrics. Mock carbohydrates (i.e., glucose, hydrolyzed guar gum, and hydrolyzed inulin) and a control (water) were modeled via a 5-fold cross-validation and hypertuned random forest model. The model’s performance improved from 64.6% accuracy when trained on only four electromagnetic features (Kappa = 0.52) to 77.2% when trained on 30 electromagnetic features (**Figure 3A**; Kappa = 0.69). Additionally, allowing 30 features resulted in a 95.6% test set accuracy (**Figure 3B**) suggesting signals captured by the VNA sufficiently discriminated among mock carbohydrates via their SCFA ratios. Furthermore, analysis of a random forest model revealed the electromagnetic features S11 and S22 unwrapped phases as the most important predictors when compared to other parameters such as resistance, reactance, and quality factor (**Figure 3C**). These phase parameters are readily associated with changes in wave delay and reflection suggesting they are central to the dielectric signature of carbohydrates.

**Figure 3.**
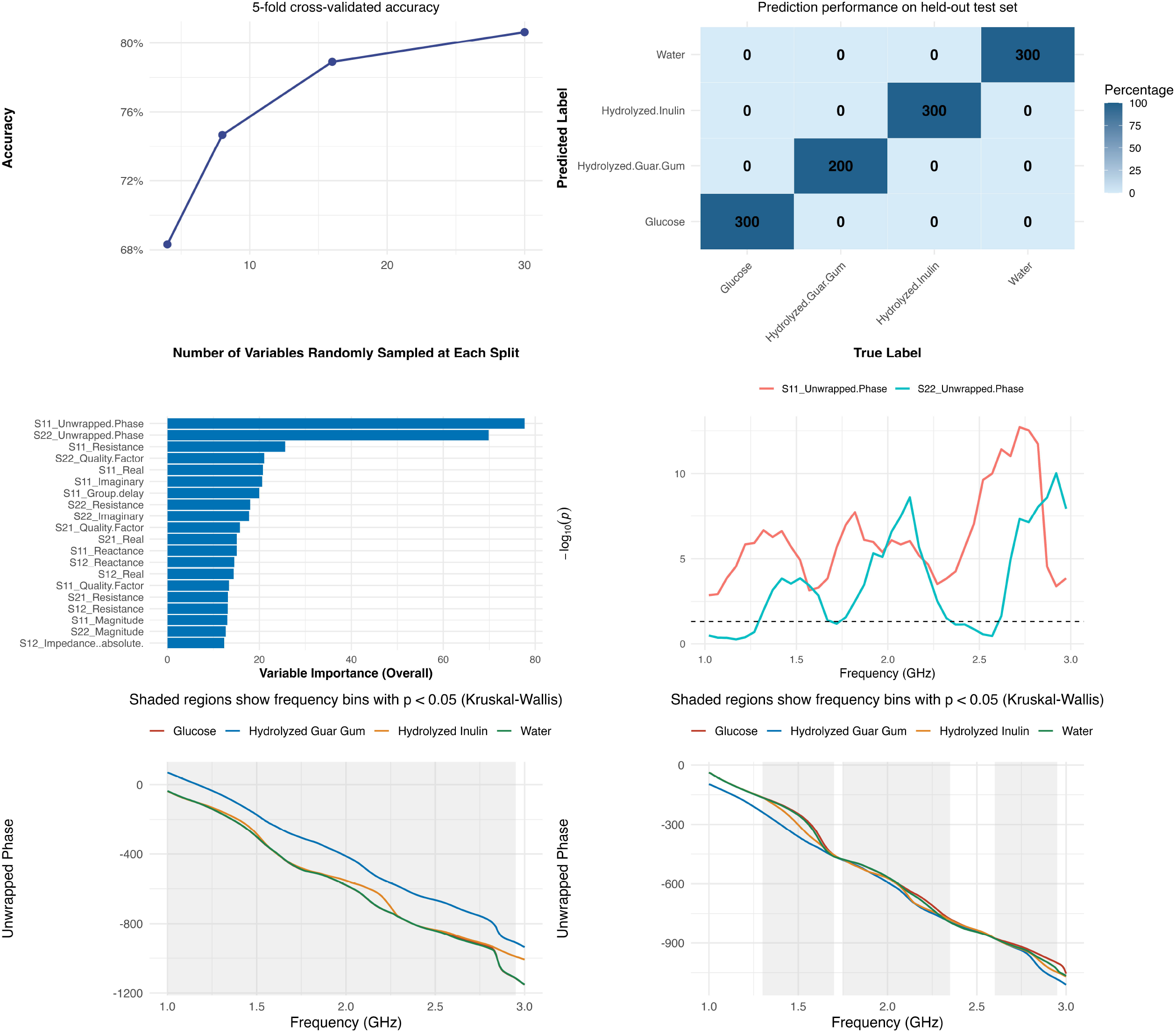
Supervised learning identifies phase features that classify carbohydrates mixtures. (**A**) Accuracy of a five-fold cross-validated random forest classifier is depicted based on a number of explanatory variables. (**B**) A confusion matrix is depicted for the four carbohydrates mixtures. (**C**) Overall variable importance (mean decrease accuracy) was reported via a cumulative bar plot for all electromagnetic properties measured by the VNA. (**D**) A per-frequency Kruskal-Wallis (50MHz bands) for S11 and S22 unwrapped phase was plotted to identify significant frequency bands (-log10p; the dashed line marks p = 0.05. (**E-F**) Class-specific unwrapped phase versus frequency for S11 (E) and S21 (F) was depicted with frequency bins as gray bands that were significant (p < 0.05).

To further analyze S11 and S22 unwrapped phase, a Kruskal-Wallis test was performed to compare frequency distributions across mock carbohydrate fermentation products, which displayed significant (p < 2.2 x 10^-16^) differences for both unwrapped phases (**Supp. Figure 3A-B**). To identify frequency bands that discriminated between mock carbohydrates, values were binned in 50 MHz intervals. A total of 28 (S11) and 28 (S22) frequency bands were significant (**Figure 3D**; p_FDR_ < 0.05). For S11, the most prominent frequency band spanned 2.5-2.85 GHz peaking at 2.72 GHz (**Figure 3E**; p_FDR_ = 1.92 x 10^-13^), while the most prominent frequency band for S22 spanned 1.4-2.5 GHz peaking at 2.92 GHz (**Figure 3D**; p_FDR_ = 9.41 x 10^-11^). Both S11 and S22 unwrapped phases displayed a broad significant frequency range suggesting both parameters possess broad zones of discriminatory power for the mock carbohydrates via consistent separability across frequencies. Combined, these results suggest the electromagnetic properties of distinct carbohydrates alter responses from VNA with frequency-resolved phase shifts as high-sensitivity markers.

## DISCUSSION

In this study, we showed acetate, propionate, and butyrate generate distinct electrical signatures that are repeatedly distinguished via their S-parameters when measured using a VNA. Importantly, SCFAs are highly distinguishable via their S21 transmission as well as their S11 and S22 unwrapped phase. Using these signals, we distinguished mixtures of SCFAs that mimic realistic fermentation outputs from varying carbohydrates. Together this points to the potential of a label-free, non-invasive approach to monitoring gut metabolic activity in real time, complementing commonly used ex vivo approaches such as stool mass spectrometry. This work is, to the best of our knowledge, among the first to leverage frequency-resolved electromagnetic properties to sense SCFAs as analytes. Our findings not only demonstrate a more practical approach to employing ingestible biosensors in order to measure SCFAs, but also establish a foundation for developing real-time, in vivo diagnostic tools capable of directly assessing gut microbial function and metabolic health throughout the GI tract.

### S-parameters distinguish among SCFAs

We identified discriminatory frequency ranges (notably ∼1.38-1.40 GHz and ∼1.79-1.85 GHz), where the S21 parameter differentiates SCFAs, with butyrate showing greater attenuation than acetate or propionate. Butyrate, being a four-carbon molecule and having less polarity, allows greater detection because of its stronger electrochemical activity and prominent electrode absorption. These band-limited differences are consistent with composition-dependent effective permittivity and loss tangent that alter how energy is transmitted and reflected, giving butyrate a stronger and more distinct signal than acetate or propionate. Importantly, the broader literature on liquid sensing has demonstrated that frequency-scale windows can reliably encode biomolecular or matrix differences, provided calibration is controlled (La Gioia et al., 2018). Our data aligns with this theory, which suggests that S-parameters convert dielectric contrasts into measurable magnitude and phase changes. Prior research has demonstrated that liquid characterization and metabolite sensing (e.g. glucose) can carry analyte information rather than a uniform broadband shift (Mimee et al. 2018). We have built on this concept by mapping frequency-resolved bands stable enough to classify SCFAs both alone and in a physiologically-relevant mixed solution. From a metabolic standpoint, these electrical signatures are valuable because SCFA production is rapid and spatially heterogeneous, and absorption by colonocytes is efficient, making fecal concentrations an improper proxy for real-time luminal flux. Real-time sensing of electrical or electromagnetic signatures, as suggested by recent advances in microwave-based metabolite detection and gut microbiome monitoring, enables the capture of transient and localized metabolic events that are otherwise missed by traditional stool analysis (Allaband et al., 2019). A method like the one we describe here, that can sample in vivo at relevant timescales, would address this long-standing gap in microbiome research.

### Machine learning inference and feature discovery

S-parameter data are highly-dimensional and correlated making supervised learning such as random forest modeling well-suited to rank informative features and integrate weak, distributed shifts across bandwidth. For example, a recent study that utilized machine learning in order to detect Alzheimer’s via microwave sensing, indicated that applying random forest modeling to S-parameter measurements had not only improved classification accuracy but also highlighted specific frequency-dependent patterns in tissue permittivity that ultimately revealed subtle biomarkers that may be indicative of early Alzheimer’s disease (Cardinali et al., 2025). In our study, S11 and S22 unwrapped phase were the top predictors of SCFA mixtures. Both S11 and S22 provide strong, distinguishable features that are able to capture how molecules interact with electromagnetic fields. Considering phase often provides higher sensitivity than magnitude when losses are modest, our results implicating S11 and S22 as prominent features aligns with their previously published use in microwave liquid sensors (Zaarour et al., 2025). S11 captures how the sample looks to the sensor at excitation, while S22 captures the response at the receiving port in the 2-port VNA. These two aspects are important because they capture complementary information. S11 reflects energy at the surface while S22 reflects how it cultivates through the sample. Together, S11 and S22 both provide a wider electromagnetic signal that strengthens material differentiation. Using both adds geometric diversity, improving robustness to nuisance factors that represent small differences. Practically, the complementarity of both phases improved classification accuracy of mock carbohydrates than either feature alone, which mirrors practices in radiofrequency liquid metrology (La Gioia et al., 2018).

### Lack of analyte and matrix realism

While we focused on acetate, propionate, and butyrate, we acknowledge that other fermentation products are pervasive throughout the gut including lactate, branched-chain fatty acids, and amino-acid derivatives (Boets et al., 2017). It is common for these metabolites to co-occur with SCFAs and co-vary across diets, presenting distinct dielectric behaviors due their differences in structure, polarity, and pKa. For example, branched-chain amino acids such as isobutyrate are able to be produced alongside SCFAs during fermentation. Their co-occurence can shift dielectric properties due to their electrochemical structure, which in result could complicate further interpretation. Additionally, our mixtures were mock solutions of fermentation products; live cultures add ionic strength, proteins, mucins, gases, and temperature drift that can affect permittivity and coupling. The dielectric-metrology literature details strategies to mitigate these confounders (temperature control, salinity baselines, probe geometry, and calibration routines) that should be adopted for in vitro and translational studies (La Gioia et al., 2018). Because we did not implement such controls, our interpretation is limited. Specifically, we cannot confidently observe permittivity changes to specific SCFAs or other metabolites, nor can we make precise, quantitative predictions about real-time luminal flux. Instead, the data reflects qualitative trends that may be influenced by multiple confounding factors, including ionic strength, proteins, pH variations, and transient microbial activity. Also, we didn’t account for the rapid SCFA metabolism and uptake, making their luminal peak transient. A successful device will require temporal resolution and spatial localization to capture true production dynamics (Boets et al., 2017). Sensing metabolites at the right time is challenging because their concentrations can fluctuate rapidly due to ongoing fermentation and microbial activity. Thus, different regions of the gut may produce distinct metabolic profiles. Because we used sodium salts at a controlled pH, future work will need to account for shifts in ionization states resulting in altered permittivity (Gabriel et al., 1996). Lastly, real diets include many polysaccharides and fibers, often in combination, producing overlapping SCFA profiles that can complicate detection and classification. In order to address this, future studies may benefit using stoichiometry of known carbohydrate fermentation to anticipate the amount of SCFAs being produced. Linking stoichiometry and electromagnetic sensing could improve accuracy within these measurements and therefore translate it into metabolic context and host diet. Our results of the mixtures suggest this is tractable, but performance should improve with multi-sensor arrays and larger, heterogeneous training sets (Zaarour et al., 2025). Further, radioisotopic or other molecular markers could be incorporated alongside microwave sensing to improve metabolite specificity and/or signal differentiation when analyzing complex biological samples. For in vivo deployment, miniaturization, biocompatible packaging, and wireless telemetry will be required, all of which are active areas in ingestible and wearable sensors.

### Conclusion

SCFAs are central to host-microbe interactions shaping metabolism, immune tone, and barrier function (Koh et al., 2016). By showing individual SCFAs and SCFA mixtures have reproducible, frequency-resolved dielectric signatures - and identifying the most discriminative features - we provide a foundation for dynamic, label-free monitoring that could be embedded in an ingestible or minimally invasive device. These systems are increasingly feasible, which have been demonstrated by recent ingestible biosensor platforms with on-board telemetry (Mimee et al., 2018). Our work underlines the need for adapting these sensors for real-time measurement of gut metabolite levels. Finally, microwave biosensing has matured in glucose monitoring of biofluids and dielectric liquid metrology, supporting the idea that solute-dependent frequency fingerprints are robust when instrumentation is controlled - an approach our results now extend to SCFAs.

## AUTHOR CONTRIBUTIONS

JHM and DV conceived and developed the theoretical framework for the study. JHM, DV, and SR contributed to study design and methodology. KP and SR performed the experiments with the assistance of DV. KP, SR, JM, DV, and JHM carried out computational analysis of the data. All authors contributed to the interpretation of the results and writing of the manuscript.

## ACKNOWLEDGEMENTS

We are grateful to the individuals who contributed to the success of this research study and manuscript. Notably, we would like to express gratitude to Trung Ha from the UIC Engineering Department for his suggestion of and assistance with using a LibreVNA. Also from the UIC Engineering department, we would like to thank Shafkat Hossain for his general advice on vector network analyzers. Finally, we acknowledge the substantial support that the UIC Biological Sciences and UIC Engineering departments provided for the continuation and completion of this research study and manuscript.

## DATA AVAILABILITY STATEMENT

The data underlying this study are openly available on Github at [https://github.com/jud-m/SC-FAs_VNA].

## ABBREVIATIONS

dH2O: Deionized water
GHz: Gigahertz
GM: Gut microbiome
M: Molar
MHz: Megahertz
mM: Millimolar
MeV: Mega-electron Volt
mV: Millivolt
SCFA: Short chain fatty acid
S21: Scattering parameter from port 1 to port 2
S11: Scattering parameter of input reflection coefficient
S22: Scattering parameter of output reflection coefficient
uM: Micromolar
VNA: Vector Network Analyzer

