## Supplemental Figures 1-3 for "Real-time measurement of short-chain fatty acids via microwave sensing: A pilot study"

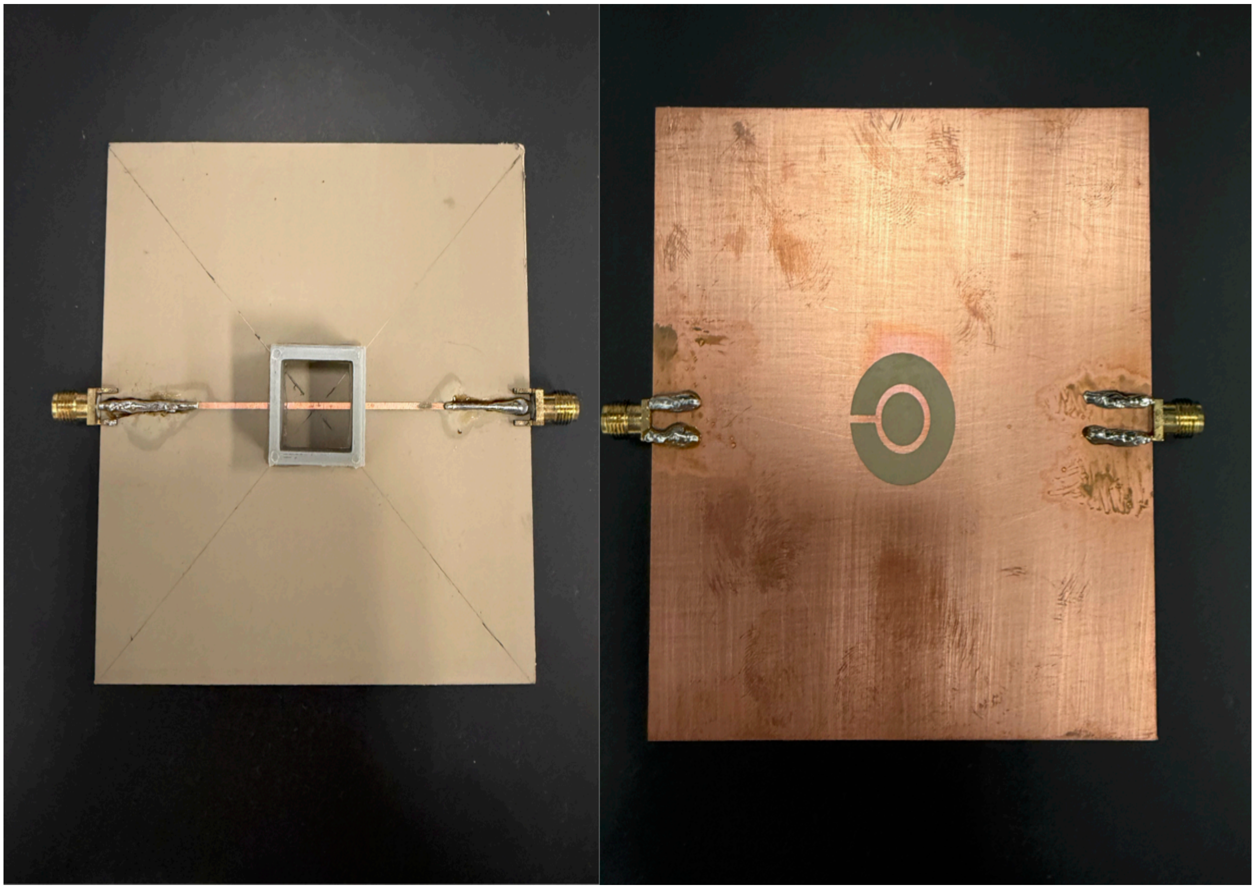

**Supplemental Figure 1.** *Overview of the sensor plate design.* The sensor plate consists of an array of integrated sensing elements arranged to ensure proper uniform signal acquisition across the measurement surface. The layout highlights the positioning of electrodes, connection ports, and structural components used during data collection with a 3D printed part attached to the center to allow the sample to pass through. This figure provides a visual reference for the hardware configuration used throughout the experiments.

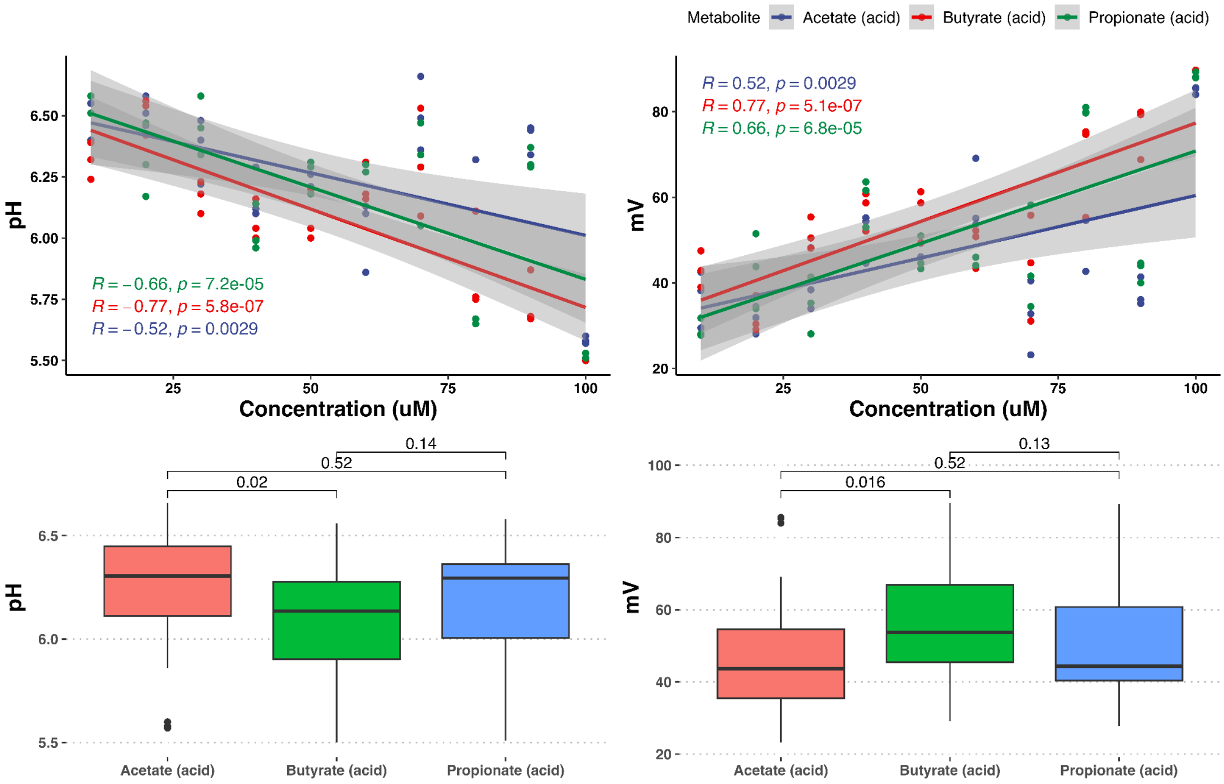


**Supplemental Figure 2**. *Electrochemical profiles of SCFAs across concentrations*. A linear model was generated to assess the correlation between either (**A**) pH and concentration or (**B**) mV and concentration (10-100 uM). A Pearson correlation was employed to determine the correlation coefficient. (C-D) Boxplots were generated to compare pH **(C)** and mV **(D)** among SCFAs. Significance was determined via Kruskal-Wallis.

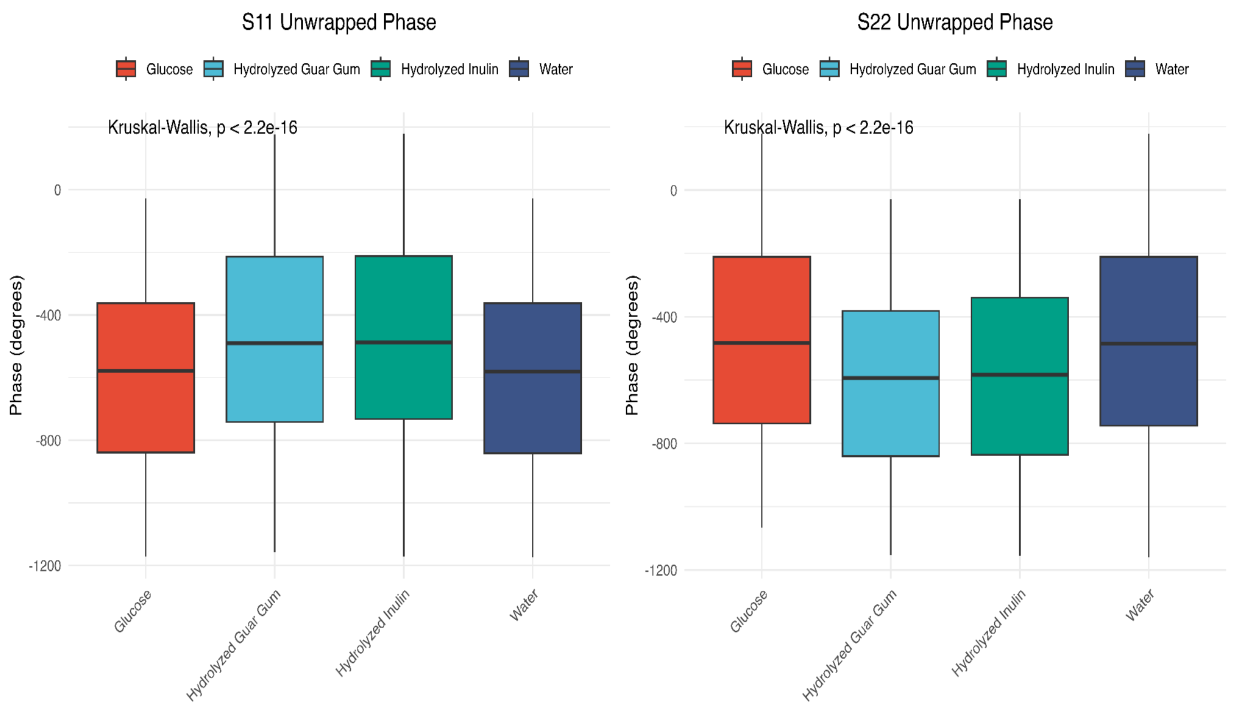


**Supplemental Figure 3.** *Phase-based reflection features differ by carbohydrate mixture*. Boxplots were generated for (A) S11 unwrapped phase and (B) S21 unwrapped phase for the four carbohydrate mixtures across frequencies 1-3 GHz. A Kruskal-Wallis was performed to determine significance.
