## Supplemental Table 1 for "Real-time measurement of short-chain fatty acids via microwave sensing: A pilot study"

**Supplemental Table 1.** *Scattering Parameter Measurement Descriptions.* S-Parameters characterize the electrical behavior of electrical linear networks when undergoing steady-state stimulation by electrical signals. S11 and S22 represent reflection coefficients at ports 1 and 2, while S21 and S12 represent transmission coefficients between the ports.

| | Parameter | Description | Unit |
|---|---|---|---|
| **General** | Frequency | is the rate at which an electrical signal oscillates or repeats a complete cycle over a specific unit of time, typically one second. | GHz |

| | Parameter | Description | Unit |
|---|---|---|---|
| **S11 Measurements (Port 1 Reflection Characteristics)** | S11_ Magnitude | Magnitude of the reflection coefficient at port 1 ; indicates the proportion of power reflected back | dB |
| | S11_ Magnitude (linear) | Linear scale magnitude of S11 ; represents the direct ratio of reflected power to incident power | – |
| | S11_ Phase | Phase angle of the S11 reflection coefficient | Degrees (°) |

| | | |
|---|---|---|
| S11_ Unwrapped Phase | Continuous phase variation of S11 without discontinuities | Degrees (°) |
| S11_ VSWR | Voltage standing wave ratio at port 1 ; a measure of impedance matching quality | – |
| S11_Real | Real (resistive) component of the S11 complex reflection coefficient | – |
| S11_Imaginary | Imaginary (reactive) component of the S11 complex reflection coefficient | – |
| S11_Impedance (absolute) | Magnitude of the impedance at port 1, derived from S11 | Ω |
| S11_Resistance | Real (resistive) component of the input impedance at port 1 | Ω |
| S11_Reactance | Imaginary (reactive) component of the input impedance at port 1 ( capacitive or inductive) | Ω |
| S11_Capacitance | Equivalent capacitance at port 1 calculated from reactance | F |
| S11_Inductance | Equivalent inductance at port 1 calculated from reactance | H |

| | | | |
|---|---|---|---|
| | S11_Quality Factor | Quality factor (Q) related to losses at the input port | – |
| | S11_Group Delay | Time delay of the signal reflection at port 1 | s |

| | Parameter | Description | Unit |
|---|---|---|---|
| **S12 Measurements (Reverse Transmission: Port 2 → Port 1)** | S12_Magnitude | Magnitude of the reverse transmission coefficient from port 2 to port 1 | dB |
| | S12_Magnitude (linear) | Linear scale magnitude of the S12 transmission coefficient | – |
| | S12_Phase | Phase angle of the S12 transmission coefficient | Degrees (°) |
| | S12_Unwrapped Phase | Continuous phase variation of S12 without discontinuities | Degrees (°) |
| | S12_VSWR | Voltage standing wave ratio for S12 (less commonly used for transmission parameters) | – |

| | Parameter | Description | Unit |
|---|---|---|---|
| | S12_Real | Real component of the S12 complex transmission coefficient | – |
| | S12_Imaginary | Imaginary component of the S12 complex transmission coefficient | – |
| | S12_Impedance (absolute) | Magnitude of impedance associated with S12 parameter | Ω |
| | S12_Resistance | Resistance component related to S12 | Ω |
| | S12_Reactance | Reactance component related to S12 | Ω |
| | S12_Capacitance | Equivalent inductance related to S12 | F |
| | S12_Inductance | Equivalent inductance related to S12 | H |
| | S12_Quality Factor | Quality factor related to S12 transmission | – |
| | S12_Group Delay | Time delay of transmission from port 2 to port 1 | s |

| | Parameter | Description | Unit |
|---|---|---|---|
| | S21_Magnitude | Magnitude of the forward transmission coefficient from port 1 to port 2 | dB |

| | | | |
|---|---|---|---|
| **S21 Measurements (Forward Transmission: Port 1 → Port 2)** | S21_Magnitude (linear) | Linear scale magnitude of the S21 transmission coefficient | – |
| | S21_Phase | Phase angle of the S21 transmission coefficient | Degrees (°) |
| | S21_Unwrapped Phase | Continuous phase variation of S21 without discontinuities | Degrees (°) |
| | S21_VSWR | Voltage standing wave ratio for S21 ( less commonly used for transmission parameters) | – |
| | S21_Real | Real component of the S21 complex transmission coefficient | – |
| | S21_Imaginary | Imaginary component of the S21 complex transmission coefficient | – |
| | S21_Impedance(absolute) | Magnitude of impedance associated with S21 parameter | Ω |
| | S21_Resistance | Resistance component related to S21 | Ω |
| | S21_Reactance | Reactance component related to S21 | Ω |
| | S21_Capacitance | Equivalent capacitance related to S21 | F |
| | S21_Inductance | Equivalent inductance related to S21 | H |
| | S21_Quality Factor | Quality factor related to S21 transmission | – |
| | S21_Group Delay | Time delay of transmission from port 1 to port 2 | s |

| | Parameter | Description | Unit |
|---|---|---|---|
| **S22 Measurements (Port 2 Reflection Characteristics)** | S22_Magnitude | Magnitude of the reflection coefficient at port 2 | dB |
| | S22_Magnitude (linear) | Linear scale magnitude of the S22 reflection coefficient | – |
| | S22_Phase | Phase angle of the S22 reflection coefficient | Degrees (°) |
| | S22_Unwrapped Phase | Continuous phase variation of S22 without discontinuities | Degrees (°) |
| | S22_VSWR | Voltage standing wave ratio at port 2 | – |
| | S22_Real | Real (resistive) component of the S22 complex reflection coefficient | – |
| | S22_Imaginary | Imaginary (reactive) component of the S22 complex transmission reflection coefficient | – |
| | S22_Impedance (absolute) | Magnitude of the impedance at port 2 , derived from S22 | Ω |

| | | | |
|---|---|---|---|
| | S22_Resistance | Real (resistive) component of the input impedance at port 2 | Ω |
| | S22_Reactance | Imaginary (reactive) component of the input impedance at port 2 | Ω |
| | S22_Capacitance | Equivalent capacitance at port 2 calculated from reactance | F |
| | S22_Inductance | Equivalent inductance at port 2 calculated from reactance | H |
| | S22_Quality Factor | Quality factor related to losses at the output port ; higher values indicate lower loss | – |
| | S22_Group Delay | Time delay of the signal reflection at port 2 | s |
