## Supplemental Table 2 for "Real-time measurement of short-chain fatty acids via microwave sensing: A pilot study"

**Supplemental Table 2**. *Pairwise contrasts among SCFAs across frequency bands*. Wilcoxon rank-sum with corrected p-values (Benjamini-Hochberg) were generated to identify frequency bands where S21 differed by SCFA.

| Frequency_scaled | Contrast | Estimate | Standard Error | Degrees of freedom | T_ratio | P-value | P-value adjusted |
|---|---|---|---|---|---|---|---|
| 1 | acetate - propionate | -0.4154557333 | 0.09691374475 | 33 | -4.2868608 | 1.48E-04 | 4.99E-04 |
| 1 | acetate - butyrate | 0.3816027917 | 0.09691374475 | 33 | 3.937550785 | 4.02E-04 | 0.001229180278 |
| 1 | propionate - butyrate | 0.797058525 | 0.09691374475 | 33 | 8.224411584 | 1.70E-09 | 1.86E-08 |
| 1.01 | acetate - propionate | -0.4079265167 | 0.1164970052 | 33 | -3.501605179 | 0.001349524768 | 0.003767423312 |
| 1.01 | acetate - butyrate | 0.3714798333 | 0.1164970052 | 33 | 3.188750069 | 0.003124390158 | 0.008085868091 |
| 1.01 | propionate - butyrate | 0.77940635 | 0.1164970052 | 33 | 6.690355248 | 1.28E-07 | 1.16E-06 |
| 1.02 | acetate - propionate | -0.3951691417 | 0.1373480377 | 33 | -2.87713715 | 0.006983922478 | 0.01677810858 |
| 1.02 | acetate - butyrate | 0.3451531583 | 0.1373480377 | 33 | 2.512982087 | 0.01703544972 | 0.03749042401 |
| 1.02 | propionate - butyrate | 0.7403223 | 0.1373480377 | 33 | 5.390119237 | 5.83E-06 | 3.65E-05 |
| 1.03 | acetate - propionate | -0.3849798 | 0.159892611 | 33 | -2.40773978 | 0.02180173954 | 0.04661861328 |
| 1.03 | acetate - butyrate | 0.3334610083 | 0.159892611 | 33 | 2.085531071 | 0.04483193217 | 0.08863493474 |
| 1.03 | propionate - butyrate | 0.7184408083 | 0.159892611 | 33 | 4.493270851 | 8.14E-05 | 3.27E-04 |
| 1.04 | acetate - propionate | -0.3604026583 | 0.1828750263 | 33 | -1.970759297 | 0.05718735472 | 0.1098215761 |
| 1.04 | acetate - butyrate | 0.32918715 | 0.1828750263 | 33 | 1.80006618 | 0.0809982674 | 0.148455791 |
| 1.04 | propionate - butyrate | 0.6895898083 | 0.1828750263 | 33 | 3.770825477 | 6.42E-04 | 0.001887289961 |

| | | | | | | | |
|---|---|---|---|---|---|---|---|
| 1.05 | acetate - propionate | -0.33605765 | 0.2085088016 | 33 | -1.611719253 | 0.1165476455 | 0.204297181 |
| 1.05 | acetate - butyrate | 0.2880055417 | 0.2085088016 | 33 | 1.381263234 | 0.1764800929 | 0.2939709835 |
| 1.05 | propionate - butyrate | 0.6240631917 | 0.2085088016 | 33 | 2.992982487 | 0.005200017048 | 0.01279840931 |
| 1.06 | acetate - propionate | -0.31195185 | 0.2342883136 | 33 | -1.331487026 | 0.1921538245 | 0.3148607504 |
| 1.06 | acetate - butyrate | 0.2778946417 | 0.2342883136 | 33 | 1.186122506 | 0.2440439667 | 0.3882810341 |
| 1.06 | propionate - butyrate | 0.5898464917 | 0.2342883136 | 33 | 2.517609532 | 0.01684965202 | 0.03721736326 |
| 1.07 | acetate - propionate | -0.2789789583 | 0.2612234396 | 33 | -1.067970618 | 0.2932843645 | 0.4488590655 |
| 1.07 | acetate - butyrate | 0.2531406583 | 0.2612234396 | 33 | 0.9690579784 | 0.3395655133 | 0.4969854478 |
| 1.07 | propionate - butyrate | 0.5321196167 | 0.2612234396 | 33 | 2.037028597 | 0.04973365073 | 0.09642891123 |
| 1.08 | acetate - propionate | -0.2466106917 | 0.2892311224 | 33 | -0.8526423076 | 0.4000038905 | 0.5445106773 |
| 1.08 | acetate - butyrate | 0.223010925 | 0.2892311224 | 33 | 0.7710474693 | 0.4461647023 | 0.5887030974 |
| 1.08 | propionate - butyrate | 0.4696216167 | 0.2892311224 | 33 | 1.623689777 | 0.1139574259 | 0.2009249351 |
| 1.09 | acetate - propionate | -0.2081326833 | 0.3184296205 | 33 | -0.6536222447 | 0.5178840991 | 0.6616188809 |
| 1.09 | acetate - butyrate | 0.2045525917 | 0.3184296205 | 33 | 0.6423792841 | 0.5250649889 | 0.6679624226 |
| 1.09 | propionate - butyrate | 0.412685275 | 0.3184296205 | 33 | 1.296001529 | 0.2039663064 | 0.3324099534 |
| 1.1 | acetate - propionate | -0.1662863333 | 0.3485134488 | 33 | -0.47713032 | 0.6364141331 | 0.7732221747 |
| 1.1 | acetate - butyrate | 0.1801092583 | 0.3485134488 | 33 | 0.5167928497 | 0.6087471937 | 0.7522019627 |
| 1.1 | propionate - butyrate | 0.3463955917 | 0.3485134488 | 33 | 0.9939231697 | 0.3274927599 | 0.4876003314 |
| 1.11 | acetate - propionate | -0.1252422 | 0.3792726033 | 33 | -0.3302168385 | 0.7433234145 | 0.8457056961 |
| 1.11 | acetate - butyrate | 0.1769845083 | 0.3792726033 | 33 | 0.466641953 | 0.6438222063 | 0.7780055919 |

| | | | | | | | |
|---|---|---|---|---|---|---|---|
| 1.11 | propionate - butyrate | 0.3022267083 | 0.3792726033 | 33 | 0.7968587915 | 0.431229117 | 0.5752901715 |
| 1.12 | acetate - propionate | -0.0777595 | 0.4126791025 | 33 | -0.1884260665 | 0.8516966484 | 0.9173516022 |
| 1.12 | acetate - butyrate | 0.1670520667 | 0.4126791025 | 33 | 0.4047989483 | 0.6882373715 | 0.8058390972 |
| 1.12 | propionate - butyrate | 0.2448115667 | 0.4126791025 | 33 | 0.5932250147 | 0.5570748186 | 0.6969214017 |
| 1.13 | acetate - propionate | -0.0354539 | 0.4466840853 | 33 | -0.07937130774 | 0.9372166612 | 0.9594934579 |
| 1.13 | acetate - butyrate | 0.149166825 | 0.4466840853 | 33 | 0.3339425556 | 0.7405358222 | 0.8441268446 |
| 1.13 | propionate - butyrate | 0.184620725 | 0.4466840853 | 33 | 0.4133138633 | 0.6820502509 | 0.8031132307 |
| 1.14 | acetate - propionate | 0.009559666667 | 0.481086667 | 33 | 0.0198709865 | 0.9842659532 | 0.9899678042 |
| 1.14 | acetate - butyrate | 0.170314925 | 0.481086667 | 33 | 0.3540212953 | 0.7255749989 | 0.8381642229 |
| 1.14 | propionate - butyrate | 0.1607552583 | 0.481086667 | 33 | 0.3341503088 | 0.7403804851 | 0.8441268446 |
| 1.15 | acetate - propionate | 0.0616722 | 0.5193140668 | 33 | 0.1187570373 | 0.9061878257 | 0.943334721 |
| 1.15 | acetate - butyrate | 0.18790845 | 0.5193140668 | 33 | 0.3618397075 | 0.7197785654 | 0.8330642513 |
| 1.15 | propionate - butyrate | 0.12623625 | 0.5193140668 | 33 | 0.2430826702 | 0.8094460232 | 0.8911838872 |
| 1.16 | acetate - propionate | 0.1154862417 | 0.5602037121 | 33 | 0.2061504399 | 0.8379399946 | 0.9087730517 |
| 1.16 | acetate - butyrate | 0.2035729417 | 0.5602037121 | 33 | 0.3633909188 | 0.7186305123 | 0.8330642513 |
| 1.16 | propionate - butyrate | 0.0880867 | 0.5602037121 | 33 | 0.1572404789 | 0.8760134719 | 0.9283587409 |
| 1.17 | acetate - propionate | 0.1650048333 | 0.6041045092 | 33 | 0.2731395492 | 0.7864480931 | 0.8749597789 |
| 1.17 | acetate - butyrate | 0.249593725 | 0.6041045092 | 33 | 0.413163155 | 0.6821595662 | 0.8031132307 |
| 1.17 | propionate - butyrate | 0.08458889167 | 0.6041045092 | 33 | 0.1400236058 | 0.8894924984 | 0.9356081308 |
| 1.18 | acetate - propionate | 0.2178624917 | 0.6517535955 | 33 | 0.334271254 | 0.7402900594 | 0.8441268446 |

| | | | | | | | |
|---|---|---|---|---|---|---|---|
| 1.18 | acetate - butyrate | 0.2883775 | 0.6517535955 | 33 | 0.4424639956 | 0.6610400817 | 0.7911493768 |
| 1.18 | propionate - butyrate | 0.07051500833 | 0.6517535955 | 33 | 0.1081927416 | 0.9144979722 | 0.9474953217 |
| 1.19 | acetate - propionate | 0.26821065 | 0.7019098114 | 33 | 0.3821155448 | 0.7048254766 | 0.8193636246 |
| 1.19 | acetate - butyrate | 0.36549225 | 0.7019098114 | 33 | 0.5207111285 | 0.6060445806 | 0.7504001686 |
| 1.19 | propionate - butyrate | 0.0972816 | 0.7019098114 | 33 | 0.1385955837 | 0.8906120515 | 0.9356081308 |
| 1.2 | acetate - propionate | 0.32146735 | 0.7573602857 | 33 | 0.4244576275 | 0.6739866671 | 0.8000274809 |
| 1.2 | acetate - butyrate | 0.4699826167 | 0.7573602857 | 33 | 0.6205535536 | 0.5391558532 | 0.6801484926 |
| 1.2 | propionate - butyrate | 0.1485152667 | 0.7573602857 | 33 | 0.1960959262 | 0.8457376739 | 0.913942325 |
| 1.21 | acetate - propionate | 0.3748649167 | 0.8208842774 | 33 | 0.4566598813 | 0.6509072393 | 0.7834272761 |
| 1.21 | acetate - butyrate | 0.58309335 | 0.8208842774 | 33 | 0.7103234476 | 0.4824935839 | 0.623005634 |
| 1.21 | propionate - butyrate | 0.2082284333 | 0.8208842774 | 33 | 0.2536635663 | 0.8013292712 | 0.8882381444 |
| 1.22 | acetate - propionate | 0.4275518667 | 0.8935902976 | 33 | 0.4784652069 | 0.6354739859 | 0.7732221747 |
| 1.22 | acetate - butyrate | 0.7356928 | 0.8935902976 | 33 | 0.823299897 | 0.4162477786 | 0.560262077 |
| 1.22 | propionate - butyrate | 0.3081409333 | 0.8935902976 | 33 | 0.3448346901 | 0.7324068353 | 0.840700554 |
| 1.23 | acetate - propionate | 0.4841474417 | 0.9774761504 | 33 | 0.4953035851 | 0.6236680514 | 0.7660997062 |
| 1.23 | acetate - butyrate | 0.9535620083 | 0.9774761504 | 33 | 0.9755348076 | 0.336392421 | 0.4969742958 |
| 1.23 | propionate - butyrate | 0.4694145667 | 0.9774761504 | 33 | 0.4802312225 | 0.6342311432 | 0.7732221747 |
| 1.24 | acetate - propionate | 0.5324220583 | 1.07536949 | 33 | 0.4951061597 | 0.6238058968 | 0.7660997062 |
| 1.24 | acetate - butyrate | 1.229309 | 1.07536949 | 33 | 1.143150342 | 0.2612018321 | 0.4080432766 |
| 1.24 | propionate - butyrate | 0.6968869417 | 1.07536949 | 33 | 0.6480441823 | 0.5214401603 | 0.6647535236 |

| | | | | | | | |
|---|---|---|---|---|---|---|---|
| 1.25 | acetate - propionate | 0.5771830667 | 1.194644456 | 33 | 0.4831421297 | 0.6321849312 | 0.7732221747 |
| 1.25 | acetate - butyrate | 1.62887345 | 1.194644456 | 33 | 1.363479688 | 0.1819614817 | 0.3014361908 |
| 1.25 | propionate - butyrate | 1.051690383 | 1.194644456 | 33 | 0.8803375583 | 0.3850429229 | 0.5375613699 |
| 1.26 | acetate - propionate | 0.6302288667 | 1.343439485 | 33 | 0.4691159323 | 0.6420714253 | 0.7774479306 |
| 1.26 | acetate - butyrate | 2.249227325 | 1.343439485 | 33 | 1.674230473 | 0.1035389812 | 0.1847159931 |
| 1.26 | propionate - butyrate | 1.618998458 | 1.343439485 | 33 | 1.205114541 | 0.2367302327 | 0.378642786 |
| 1.27 | acetate - propionate | 0.6805189417 | 1.540528776 | 33 | 0.4417437392 | 0.6615559486 | 0.7911493768 |
| 1.27 | acetate - butyrate | 3.208215833 | 1.540528776 | 33 | 2.082541972 | 0.04512122032 | 0.08891534593 |
| 1.27 | propionate - butyrate | 2.527696892 | 1.540528776 | 33 | 1.640798232 | 0.110337912 | 0.1951136684 |
| 1.28 | acetate - propionate | 0.7299018417 | 1.826394704 | 33 | 0.3996408005 | 0.6919960696 | 0.8086698256 |
| 1.28 | acetate - butyrate | 4.974646867 | 1.826394704 | 33 | 2.723752349 | 0.0102374454 | 0.02374299836 |
| 1.28 | propionate - butyrate | 4.244745025 | 1.826394704 | 33 | 2.324111549 | 0.02642219426 | 0.05513004546 |
| 1.29 | acetate - propionate | 0.7942891667 | 2.311854705 | 33 | 0.3435722691 | 0.7333474153 | 0.840700554 |
| 1.29 | acetate - butyrate | 8.814883933 | 2.311854705 | 33 | 3.81290568 | 5.70E-04 | 0.001702594781 |
| 1.29 | propionate - butyrate | 8.020594767 | 2.311854705 | 33 | 3.46933341 | 0.001473517878 | 0.004075831561 |
| 1.3 | acetate - propionate | 0.86215685 | 2.677238157 | 33 | 0.3220321837 | 0.749459522 | 0.8473517595 |
| 1.3 | acetate - butyrate | 7.953809233 | 2.677238157 | 33 | 2.970900894 | 0.005502876391 | 0.01348875798 |
| 1.3 | propionate - butyrate | 7.091652383 | 2.677238157 | 33 | 2.64886871 | 0.01229529303 | 0.0278724124 |
| 1.31 | acetate - propionate | 0.8368912417 | 2.788763934 | 33 | 0.3000939705 | 0.7659877942 | 0.8569399627 |
| 1.31 | acetate - butyrate | -5.3590225 | 2.788763934 | 33 | -1.921647951 | 0.06332118705 | 0.1196949084 |

| | | | | | | | |
|---|---|---|---|---|---|---|---|
| 1.31 | propionate - butyrate | -6.195913742 | 2.788763934 | 33 | -2.221741922 | 0.03327189269 | 0.06778024086 |
| 1.32 | acetate - propionate | -0.8605238 | 3.139575338 | 33 | -0.2740892341 | 0.7857244914 | 0.8749597789 |
| 1.32 | acetate - butyrate | -15.92634707 | 3.139575338 | 33 | -5.072771108 | 1.49E-05 | 7.74E-05 |
| 1.32 | propionate - butyrate | -15.06582327 | 3.139575338 | 33 | -4.798681873 | 3.33E-05 | 1.51E-04 |
| 1.33 | acetate - propionate | 0.3535585333 | 2.188387644 | 33 | 0.1615611998 | 0.8726365276 | 0.9280420214 |
| 1.33 | acetate - butyrate | -9.92321025 | 2.188387644 | 33 | -4.534484682 | 7.22E-05 | 2.94E-04 |
| 1.33 | propionate - butyrate | -10.27676878 | 2.188387644 | 33 | -4.696045882 | 4.50E-05 | 1.98E-04 |
| 1.34 | acetate - propionate | 0.5121449167 | 1.596503852 | 33 | 0.320791532 | 0.7503911104 | 0.8473517595 |
| 1.34 | acetate - butyrate | -7.328848908 | 1.596503852 | 33 | -4.590561368 | 6.13E-05 | 2.53E-04 |
| 1.34 | propionate - butyrate | -7.840993825 | 1.596503852 | 33 | -4.9113529 | 2.39E-05 | 1.14E-04 |
| 1.35 | acetate - propionate | 0.565025075 | 1.187340699 | 33 | 0.475874427 | 0.6372992053 | 0.7732221747 |
| 1.35 | acetate - butyrate | -5.991706008 | 1.187340699 | 33 | -5.046324119 | 1.61E-05 | 8.15E-05 |
| 1.35 | propionate - butyrate | -6.556731083 | 1.187340699 | 33 | -5.522198546 | 3.95E-06 | 2.77E-05 |
| 1.36 | acetate - propionate | 0.5828325417 | 0.8668633608 | 33 | 0.6723464943 | 0.5060438287 | 0.6483354266 |
| 1.36 | acetate - butyrate | -5.137749533 | 0.8668633608 | 33 | -5.926827417 | 1.20E-06 | 9.49E-06 |
| 1.36 | propionate - butyrate | -5.720582075 | 0.8668633608 | 33 | -6.599173911 | 1.67E-07 | 1.48E-06 |
| 1.37 | acetate - propionate | 0.5843583833 | 0.6011903878 | 33 | 0.9720022063 | 0.3381206123 | 0.4969854478 |
| 1.37 | acetate - butyrate | -4.5536145 | 0.6011903878 | 33 | -7.574330183 | 1.03E-08 | 1.03E-07 |
| 1.37 | propionate - butyrate | -5.137972883 | 0.6011903878 | 33 | -8.54633239 | 7.08E-10 | 8.06E-09 |
| 1.38 | acetate - propionate | 0.5826309167 | 0.3899073828 | 33 | 1.494280289 | 0.1446038813 | 0.2442468919 |

| | | | | | | | |
|---|---|---|---|---|---|---|---|
| 1.38 | acetate - butyrate | -4.086415142 | 0.3899073828 | 33 | -10.48047645 | 4.93E-12 | 1.49E-10 |
| 1.38 | propionate - butyrate | -4.669046058 | 0.3899073828 | 33 | -11.97475673 | 1.48E-13 | 6.86E-12 |
| 1.39 | acetate - propionate | 0.5673079167 | 0.2756413618 | 33 | 2.058137839 | 0.04754527424 | 0.09308376742 |
| 1.39 | acetate - butyrate | -3.743141383 | 0.2756413618 | 33 | -13.57975218 | 4.61E-15 | 3.97E-13 |
| 1.39 | propionate - butyrate | -4.3104493 | 0.2756413618 | 33 | -15.63789002 | 8.05E-17 | 2.88E-14 |
| 1.4 | acetate - propionate | 0.54025825 | 0.3274855337 | 33 | 1.649716383 | 0.1084891374 | 0.192408676 |
| 1.4 | acetate - butyrate | -3.425843025 | 0.3274855337 | 33 | -10.46105147 | 5.17E-12 | 1.49E-10 |
| 1.4 | propionate - butyrate | -3.966101275 | 0.3274855337 | 33 | -12.11076786 | 1.09E-13 | 5.94E-12 |
| 1.41 | acetate - propionate | 0.5063311667 | 0.4883137496 | 33 | 1.036897214 | 0.3073223812 | 0.4639280846 |
| 1.41 | acetate - butyrate | -3.143662483 | 0.4883137496 | 33 | -6.437792272 | 2.68E-07 | 2.24E-06 |
| 1.41 | propionate - butyrate | -3.64999365 | 0.4883137496 | 33 | -7.474689486 | 1.36E-08 | 1.35E-07 |
| 1.42 | acetate - propionate | 0.4620623 | 0.6878364558 | 33 | 0.6717618645 | 0.5064112536 | 0.6483354266 |
| 1.42 | acetate - butyrate | -2.845936017 | 0.6878364558 | 33 | -4.137518436 | 2.27E-04 | 7.33E-04 |
| 1.42 | propionate - butyrate | -3.307998317 | 0.6878364558 | 33 | -4.8092803 | 3.23E-05 | 1.49E-04 |
| 1.43 | acetate - propionate | 0.4071249667 | 0.9051517509 | 33 | 0.4497864212 | 0.6558051572 | 0.7877500195 |
| 1.43 | acetate - butyrate | -2.58166605 | 0.9051517509 | 33 | -2.852191411 | 0.007436881926 | 0.0177250585 |
| 1.43 | propionate - butyrate | -2.988791017 | 0.9051517509 | 33 | -3.301977833 | 0.002313571387 | 0.006172936046 |
| 1.44 | acetate - propionate | 0.3440372667 | 1.143695042 | 33 | 0.3008120643 | 0.7654449653 | 0.8569399627 |
| 1.44 | acetate - butyrate | -2.335748583 | 1.143695042 | 33 | -2.042282686 | 0.04918086029 | 0.09566470566 |
| 1.44 | propionate - butyrate | -2.67978585 | 1.143695042 | 33 | -2.34309475 | 0.02530180821 | 0.05316024512 |

| | | | | | | | |
|---|---|---|---|---|---|---|---|
| 1.45 | acetate - propionate | 0.2541584 | 1.408959539 | 33 | 0.180387295 | 0.857951757 | 0.9219882131 |
| 1.45 | acetate - butyrate | -2.113330283 | 1.408959539 | 33 | -1.49992262 | 0.143141834 | 0.2431395097 |
| 1.45 | propionate - butyrate | -2.367488683 | 1.408959539 | 33 | -1.680309915 | 0.1023408293 | 0.1831202376 |
| 1.46 | acetate - propionate | 0.1490728833 | 1.716768427 | 33 | 0.08683342551 | 0.9313286772 | 0.9580460557 |
| 1.46 | acetate - butyrate | -1.915865067 | 1.716768427 | 33 | -1.11597175 | 0.2724947557 | 0.423490561 |
| 1.46 | propionate - butyrate | -2.06493795 | 1.716768427 | 33 | -1.202805176 | 0.2376108073 | 0.3790458117 |
| 1.47 | acetate - propionate | 0.00739 | 2.096943237 | 33 | 0.003524177417 | 0.9972093377 | 0.9972093377 |
| 1.47 | acetate - butyrate | -1.737758833 | 2.096943237 | 33 | -0.8287104785 | 0.4132222237 | 0.5599393278 |
| 1.47 | propionate - butyrate | -1.745148833 | 2.096943237 | 33 | -0.8322346559 | 0.4112588768 | 0.5585340151 |
| 1.48 | acetate - propionate | -0.2141189167 | 2.614312011 | 33 | -0.08190258689 | 0.9352189359 | 0.9590765618 |
| 1.48 | acetate - butyrate | -1.596561692 | 2.614312011 | 33 | -0.610700515 | 0.545581411 | 0.6868175174 |
| 1.48 | propionate - butyrate | -1.382442775 | 2.614312011 | 33 | -0.5287979281 | 0.6004845821 | 0.7465818619 |
| 1.49 | acetate - propionate | -0.4193759833 | 3.304011303 | 33 | -0.1269293428 | 0.8997665924 | 0.9419431514 |
| 1.49 | acetate - butyrate | -1.418814092 | 3.304011303 | 33 | -0.4294216821 | 0.6704072 | 0.7973482082 |
| 1.49 | propionate - butyrate | -0.9994381083 | 3.304011303 | 33 | -0.3024923393 | 0.7641752659 | 0.8569399627 |
| 1.5 | acetate - propionate | 0.417921975 | 3.413888129 | 33 | 0.122418181 | 0.9033103319 | 0.9429222118 |
| 1.5 | acetate - butyrate | -0.7651455667 | 3.413888129 | 33 | -0.2241273111 | 0.8240401182 | 0.8995675192 |
| 1.5 | propionate - butyrate | -1.183067542 | 3.413888129 | 33 | -0.3465454921 | 0.7311328551 | 0.840700554 |
| 1.51 | acetate - propionate | 0.51938755 | 2.907172253 | 33 | 0.1786573016 | 0.8592991306 | 0.9219882131 |
| 1.51 | acetate - butyrate | -0.677397675 | 2.907172253 | 33 | -0.2330091291 | 0.8171935665 | 0.8949832031 |

| | | | | | | | |
|---|---|---|---|---|---|---|---|
| 1.51 | propionate - butyrate | -1.196785225 | 2.907172253 | 33 | -0.4116664307 | 0.6832455843 | 0.8031132307 |
| 1.52 | acetate - propionate | 0.2438530833 | 2.417694368 | 33 | 0.1008618321 | 0.9202704635 | 0.9502107697 |
| 1.52 | acetate - butyrate | -0.5940073667 | 2.417694368 | 33 | -0.24569167 | 0.8074425926 | 0.8911838872 |
| 1.52 | propionate - butyrate | -0.83786045 | 2.417694368 | 33 | -0.346553502 | 0.7311268922 | 0.840700554 |
| 1.53 | acetate - propionate | 0.05781305 | 2.094735906 | 33 | 0.02759920706 | 0.978148047 | 0.9863265424 |
| 1.53 | acetate - butyrate | -0.49808005 | 2.094735906 | 33 | -0.2377770146 | 0.813524213 | 0.893542988 |
| 1.53 | propionate - butyrate | -0.5558931 | 2.094735906 | 33 | -0.2653762216 | 0.7923704549 | 0.8799252012 |
| 1.54 | acetate - propionate | -0.09279784167 | 1.854527279 | 33 | -0.05003854229 | 0.9603933224 | 0.9733061738 |
| 1.54 | acetate - butyrate | -0.401571875 | 1.854527279 | 33 | -0.2165359764 | 0.8299030451 | 0.9016784436 |
| 1.54 | propionate - butyrate | -0.3087740333 | 1.854527279 | 33 | -0.1664974341 | 0.8687814892 | 0.9255746254 |
| 1.55 | acetate - propionate | -0.2210728583 | 1.665181551 | 33 | -0.1327620152 | 0.8951878485 | 0.9387796046 |
| 1.55 | acetate - butyrate | -0.32688625 | 1.665181551 | 33 | -0.1963066729 | 0.8455740661 | 0.913942325 |
| 1.55 | propionate - butyrate | -0.1058133917 | 1.665181551 | 33 | -0.06354465771 | 0.9497161569 | 0.9701117992 |
| 1.56 | acetate - propionate | -0.33482955 | 1.508625822 | 33 | -0.2219434039 | 0.8257257537 | 0.8995675192 |
| 1.56 | acetate - butyrate | -0.2585983833 | 1.508625822 | 33 | -0.1714132024 | 0.8649456739 | 0.9247557471 |
| 1.56 | propionate - butyrate | 0.07623116667 | 1.508625822 | 33 | 0.05053020145 | 0.9600045021 | 0.9733061738 |
| 1.57 | acetate - propionate | -0.434314675 | 1.377055093 | 33 | -0.3153938264 | 0.7544485905 | 0.8487546644 |
| 1.57 | acetate - butyrate | -0.1964365583 | 1.377055093 | 33 | -0.1426497453 | 0.8874342382 | 0.9356081308 |
| 1.57 | propionate - butyrate | 0.2378781167 | 1.377055093 | 33 | 0.1727440811 | 0.8639077463 | 0.9247557471 |
| 1.58 | acetate - propionate | -0.5199183083 | 1.261262443 | 33 | -0.4122205581 | 0.6828434312 | 0.8031132307 |

| | | | | | | | |
|---|---|---|---|---|---|---|---|
| 1.58 | acetate - butyrate | -0.1454046667 | 1.261262443 | 33 | -0.1152850205 | 0.9089178655 | 0.943334721 |
| 1.58 | propionate - butyrate | 0.3745136417 | 1.261262443 | 33 | 0.2969355376 | 0.7683767725 | 0.8580207293 |
| 1.59 | acetate - propionate | -0.6079538417 | 1.156894296 | 33 | -0.5255050905 | 0.6027456291 | 0.7478510584 |
| 1.59 | acetate - butyrate | -0.1006058833 | 1.156894296 | 33 | -0.08696203594 | 0.9312272306 | 0.9580460557 |
| 1.59 | propionate - butyrate | 0.5073479583 | 1.156894296 | 33 | 0.4385430546 | 0.6638503957 | 0.7911493768 |
| 1.6 | acetate - propionate | -0.6793871583 | 1.062903673 | 33 | -0.6391803654 | 0.5271178432 | 0.6691622304 |
| 1.6 | acetate - butyrate | -0.06391223333 | 1.062903673 | 33 | -0.06012984519 | 0.9524149007 | 0.9701117992 |
| 1.6 | propionate - butyrate | 0.615474925 | 1.062903673 | 33 | 0.5790505202 | 0.5664869897 | 0.7072290989 |
| 1.61 | acetate - propionate | -0.74686115 | 0.9762936477 | 33 | -0.7649964248 | 0.4497102126 | 0.5907957695 |
| 1.61 | acetate - butyrate | -0.03294808333 | 0.9762936477 | 33 | -0.03374812835 | 0.9732813065 | 0.9847124628 |
| 1.61 | propionate - butyrate | 0.7139130667 | 0.9762936477 | 33 | 0.7312482964 | 0.4697883969 | 0.611841044 |
| 1.62 | acetate - propionate | -0.7984958417 | 0.8972937714 | 33 | -0.8898934408 | 0.379964902 | 0.5375613699 |
| 1.62 | acetate - butyrate | -0.01180163333 | 0.8972937714 | 33 | -0.01315247437 | 0.9895853419 | 0.9928784711 |
| 1.62 | propionate - butyrate | 0.7866942083 | 0.8972937714 | 33 | 0.8767409664 | 0.3869653512 | 0.5376249795 |
| 1.63 | acetate - propionate | -0.8490400417 | 0.8239347293 | 33 | -1.030470026 | 0.3102831669 | 0.4665854106 |
| 1.63 | acetate - butyrate | 0.007686016667 | 0.8239347293 | 33 | 0.009328429054 | 0.9926132657 | 0.9942621249 |
| 1.63 | propionate - butyrate | 0.8567260583 | 0.8239347293 | 33 | 1.039798455 | 0.3059923082 | 0.4636014117 |
| 1.64 | acetate - propionate | -0.8938970083 | 0.7547298 | 33 | -1.184393419 | 0.2447179868 | 0.3883288054 |
| 1.64 | acetate - butyrate | 0.02320708333 | 0.7547298 | 33 | 0.03074886315 | 0.975655043 | 0.9854606213 |
| 1.64 | propionate - butyrate | 0.9171040917 | 0.7547298 | 33 | 1.215142282 | 0.2329345759 | 0.3735626311 |

| | | | | | | | |
|---|---|---|---|---|---|---|---|
| 1.65 | acetate - propionate | -0.9243404167 | 0.6905304287 | 33 | -1.338594765 | 0.1898522697 | 0.3119371081 |
| 1.65 | acetate - butyrate | 0.04269635 | 0.6905304287 | 33 | 0.06183123614 | 0.9510702093 | 0.9701117992 |
| 1.65 | propionate - butyrate | 0.9670367667 | 0.6905304287 | 33 | 1.400426001 | 0.1707185891 | 0.2859536367 |
| 1.66 | acetate - propionate | -0.9519272583 | 0.6292699771 | 33 | -1.51274857 | 0.1398624864 | 0.2386514157 |
| 1.66 | acetate - butyrate | 0.053607375 | 0.6292699771 | 33 | 0.0851897865 | 0.9326252649 | 0.9580460557 |
| 1.66 | propionate - butyrate | 1.005534633 | 0.6292699771 | 33 | 1.597938357 | 0.1195892871 | 0.2084171681 |
| 1.67 | acetate - propionate | -0.9763874583 | 0.5697849609 | 33 | -1.713606931 | 0.09598158342 | 0.1727668502 |
| 1.67 | acetate - butyrate | 0.06601746667 | 0.5697849609 | 33 | 0.1158638279 | 0.9084626713 | 0.943334721 |
| 1.67 | propionate - butyrate | 1.042404925 | 0.5697849609 | 33 | 1.829470759 | 0.07637691997 | 0.1425860147 |
| 1.68 | acetate - propionate | -0.9939152167 | 0.5146065429 | 33 | -1.931408044 | 0.06205875686 | 0.117677454 |
| 1.68 | acetate - butyrate | 0.07171445 | 0.5146065429 | 33 | 0.1393578278 | 0.8900144322 | 0.9356081308 |
| 1.68 | propionate - butyrate | 1.065629667 | 0.5146065429 | 33 | 2.070765872 | 0.04627699686 | 0.09089586028 |
| 1.69 | acetate - propionate | -1.006453625 | 0.4624642871 | 33 | -2.176283992 | 0.03679394758 | 0.07395583463 |
| 1.69 | acetate - butyrate | 0.0777088 | 0.4624642871 | 33 | 0.168032002 | 0.8675837015 | 0.9255746254 |
| 1.69 | propionate - butyrate | 1.084162425 | 0.4624642871 | 33 | 2.344315994 | 0.02523122382 | 0.05316024512 |
| 1.7 | acetate - propionate | -1.0186133 | 0.4115783513 | 33 | -2.474895234 | 0.01863755656 | 0.04057200941 |
| 1.7 | acetate - butyrate | 0.07742584167 | 0.4115783513 | 33 | 0.1881193251 | 0.851935153 | 0.9173516022 |
| 1.7 | propionate - butyrate | 1.096039142 | 0.4115783513 | 33 | 2.663014559 | 0.01187933657 | 0.02703109415 |
| 1.71 | acetate - propionate | -1.020882383 | 0.3641778408 | 33 | -2.803252337 | 0.008406615356 | 0.01987917278 |
| 1.71 | acetate - butyrate | 0.080476625 | 0.3641778408 | 33 | 0.2209816633 | 0.8264683344 | 0.8995675192 |

| | | | | | | | |
|---|---|---|---|---|---|---|---|
| 1.71 | propionate - butyrate | 1.101359008 | 0.3641778408 | 33 | 3.024234 | 0.004798198207 | 0.01190663999 |
| 1.72 | acetate - propionate | -1.022295242 | 0.319045979 | 33 | -3.204225437 | 0.002999411937 | 0.007829633757 |
| 1.72 | acetate - butyrate | 0.07983196667 | 0.319045979 | 33 | 0.2502208832 | 0.803967788 | 0.8895276627 |
| 1.72 | propionate - butyrate | 1.102127208 | 0.319045979 | 33 | 3.45444632 | 0.001534348778 | 0.004205510515 |
| 1.73 | acetate - propionate | -1.018745717 | 0.2756536353 | 33 | -3.695745625 | 7.91E-04 | 0.002292854687 |
| 1.73 | acetate - butyrate | 0.08750459167 | 0.2756536353 | 33 | 0.3174439966 | 0.7529066259 | 0.848603169 |
| 1.73 | propionate - butyrate | 1.106250308 | 0.2756536353 | 33 | 4.013189622 | 3.24E-04 | 0.001007342243 |
| 1.74 | acetate - propionate | -1.013656367 | 0.2361219786 | 33 | -4.292935256 | 1.45E-04 | 4.93E-04 |
| 1.74 | acetate - butyrate | 0.09009798333 | 0.2361219786 | 33 | 0.3815738961 | 0.7052234182 | 0.8193636246 |
| 1.74 | propionate - butyrate | 1.10375435 | 0.2361219786 | 33 | 4.674509153 | 4.80E-05 | 2.08E-04 |
| 1.75 | acetate - propionate | -1.007199808 | 0.2014431767 | 33 | -4.999920199 | 1.85E-05 | 9.20E-05 |
| 1.75 | acetate - butyrate | 0.09672431667 | 0.2014431767 | 33 | 0.4801568276 | 0.6342834773 | 0.7732221747 |
| 1.75 | propionate - butyrate | 1.103924125 | 0.2014431767 | 33 | 5.480077027 | 4.47E-06 | 2.96E-05 |
| 1.76 | acetate - propionate | -0.9963867 | 0.1675625018 | 33 | -5.946358458 | 1.13E-06 | 9.08E-06 |
| 1.76 | acetate - butyrate | 0.10611245 | 0.1675625018 | 33 | 0.6332708622 | 0.5309214087 | 0.6711647997 |
| 1.76 | propionate - butyrate | 1.10249915 | 0.1675625018 | 33 | 6.579629321 | 1.77E-07 | 1.55E-06 |
| 1.77 | acetate - propionate | -0.987413575 | 0.1368923172 | 33 | -7.213067868 | 2.86E-08 | 2.74E-07 |
| 1.77 | acetate - butyrate | 0.1071028083 | 0.1368923172 | 33 | 0.7823872843 | 0.4395653397 | 0.58463009 |
| 1.77 | propionate - butyrate | 1.094516383 | 0.1368923172 | 33 | 7.995455153 | 3.18E-09 | 3.37E-08 |
| 1.78 | acetate - propionate | -0.9672283833 | 0.1122742829 | 33 | -8.614870287 | 5.89E-10 | 6.83E-09 |

| | | | | | | | |
|---|---|---|---|---|---|---|---|
| 1.78 | acetate - butyrate | 0.1163131917 | 0.1122742829 | 33 | 1.035973588 | 0.3077466565 | 0.4639280846 |
| 1.78 | propionate - butyrate | 1.083541575 | 0.1122742829 | 33 | 9.650843875 | 3.91E-11 | 5.76E-10 |
| 1.79 | acetate - propionate | -0.9485390417 | 0.09136479207 | 33 | -10.38188804 | 6.28E-12 | 1.72E-10 |
| 1.79 | acetate - butyrate | 0.1243227417 | 0.09136479207 | 33 | 1.360729214 | 0.1828209287 | 0.3020301917 |
| 1.79 | propionate - butyrate | 1.072861783 | 0.09136479207 | 33 | 11.74261725 | 2.50E-13 | 9.44E-12 |
| 1.8 | acetate - propionate | -0.9224016167 | 0.0782389463 | 33 | -11.78955328 | 2.25E-13 | 9.05E-12 |
| 1.8 | acetate - butyrate | 0.142188325 | 0.0782389463 | 33 | 1.817361118 | 0.07825217434 | 0.1447425188 |
| 1.8 | propionate - butyrate | 1.064589942 | 0.0782389463 | 33 | 13.6069144 | 4.35E-15 | 3.97E-13 |
| 1.81 | acetate - propionate | -0.8928950667 | 0.0697148 7181 | 33 | -12.80781336 | 2.36E-14 | 1.42E-12 |
| 1.81 | acetate - butyrate | 0.1591913833 | 0.06971487181 | 33 | 2.28346376 | 0.02897323517 | 0.05983171509 |
| 1.81 | propionate - butyrate | 1.05208645 | 0.06971487181 | 33 | 15.09127712 | 2.26E-16 | 3.95E-14 |
| 1.82 | acetate - propionate | -0.8641304333 | 0.06715603019 | 33 | -12.8675032 | 2.07E-14 | 1.42E-12 |
| 1.82 | acetate - butyrate | 0.1799377167 | 0.06715603019 | 33 | 2.679397757 | 0.01141391145 | 0.02616953841 |
| 1.82 | propionate - butyrate | 1.04406815 | 0.06715603019 | 33 | 15.54690096 | 9.54E-17 | 2.88E-14 |
| 1.83 | acetate - propionate | -0.8348184833 | 0.06913965178 | 33 | -12.07438079 | 1.18E-13 | 5.94E-12 |
| 1.83 | acetate - butyrate | 0.2033362083 | 0.06913965178 | 33 | 2.940949269 | 0.005940344232 | 0.01450213592 |
| 1.83 | propionate - butyrate | 1.038154692 | 0.06913965178 | 33 | 15.01533006 | 2.62E-16 | 3.95E-14 |
| 1.84 | acetate - propionate | -0.8021195917 | 0.07425476962 | 33 | -10.80226355 | 2.26E-12 | 7.58E-11 |
| 1.84 | acetate - butyrate | 0.2267358083 | 0.07425476962 | 33 | 3.053484772 | 0.004448867907 | 0.01113139978 |
| 1.84 | propionate - butyrate | 1.0288554 | 0.07425476962 | 33 | 13.85574833 | 2.61E-15 | 3.15E-13 |

| | | | | | | | |
|---|---|---|---|---|---|---|---|
| 1.85 | acetate - propionate | -0.774612025 | 0.08026702163 | 33 | -9.650439362 | 3.92E-11 | 5.76E-10 |
| 1.85 | acetate - butyrate | 0.25545995 | 0.08026702163 | 33 | 3.182626499 | 0.00317520147 | 0.008182249942 |
| 1.85 | propionate - butyrate | 1.030071975 | 0.08026702163 | 33 | 12.83306586 | 2.23E-14 | 1.42E-12 |
| 1.86 | acetate - propionate | -0.7478447 | 0.08598966585 | 33 | -8.696913665 | 4.73E-10 | 5.70E-09 |
| 1.86 | acetate - butyrate | 0.2712693083 | 0.08598966585 | 33 | 3.154673363 | 0.003417311844 | 0.008768676775 |
| 1.86 | propionate - butyrate | 1.019114008 | 0.08598966585 | 33 | 11.85158703 | 1.95E-13 | 8.42E-12 |
| 1.87 | acetate - propionate | -0.7189335167 | 0.09143162104 | 33 | -7.863073065 | 4.59E-09 | 4.77E-08 |
| 1.87 | acetate - butyrate | 0.300286675 | 0.09143162104 | 33 | 3.28427596 | 0.002425477345 | 0.006386737289 |
| 1.87 | propionate - butyrate | 1.019220192 | 0.09143162104 | 33 | 11.14734902 | 9.96E-13 | 3.53E-11 |
| 1.88 | acetate - propionate | -0.691743675 | 0.09760543951 | 33 | -7.087142668 | 4.10E-08 | 3.86E-07 |
| 1.88 | acetate - butyrate | 0.3313603 | 0.09760543951 | 33 | 3.394895834 | 0.001802742138 | 0.004896637429 |
| 1.88 | propionate - butyrate | 1.023103975 | 0.09760543951 | 33 | 10.4820385 | 4.92E-12 | 1.49E-10 |
| 1.89 | acetate - propionate | -0.6652560917 | 0.1019853669 | 33 | -6.523054357 | 2.09E-07 | 1.80E-06 |
| 1.89 | acetate - butyrate | 0.3638764167 | 0.1019853669 | 33 | 3.567927712 | 0.001125516631 | 0.003171432375 |
| 1.89 | propionate - butyrate | 1.029132508 | 0.1019853669 | 33 | 10.09098207 | 1.29E-11 | 3.38E-10 |
| 1.9 | acetate - propionate | -0.6424139833 | 0.105281899 | 33 | -6.101846464 | 7.15E-07 | 5.83E-06 |
| 1.9 | acetate - butyrate | 0.398742975 | 0.105281899 | 33 | 3.787383953 | 6.13E-04 | 0.00181076686 |
| 1.9 | propionate - butyrate | 1.041156958 | 0.105281899 | 33 | 9.889230417 | 2.14E-11 | 4.17E-10 |
| 1.91 | acetate - propionate | -0.620071875 | 0.1062762012 | 33 | -5.834531795 | 1.57E-06 | 1.21E-05 |
| 1.91 | acetate - butyrate | 0.4290531833 | 0.1062762012 | 33 | 4.037152048 | 3.03E-04 | 9.56E-04 |

| | | | | | | | |
|---|---|---|---|---|---|---|---|
| 1.91 | propionate - butyrate | 1.049125058 | 0.1062762012 | 33 | 9.871683842 | 2.23E-11 | 4.17E-10 |
| 1.92 | acetate - propionate | -0.5999945583 | 0.108878747 | 33 | -5.510667369 | 4.08E-06 | 2.78E-05 |
| 1.92 | acetate - butyrate | 0.4610320167 | 0.108878747 | 33 | 4.234361888 | 1.72E-04 | 5.61E-04 |
| 1.92 | propionate - butyrate | 1.061026575 | 0.108878747 | 33 | 9.745029257 | 3.08E-11 | 5.16E-10 |
| 1.93 | acetate - propionate | -0.5837102917 | 0.1104327124 | 33 | -5.285664719 | 7.94E-06 | 4.56E-05 |
| 1.93 | acetate - butyrate | 0.49423165 | 0.1104327124 | 33 | 4.475409861 | 8.57E-05 | 3.36E-04 |
| 1.93 | propionate - butyrate | 1.077941942 | 0.1104327124 | 33 | 9.761074581 | 2.96E-11 | 5.09E-10 |
| 1.94 | acetate - propionate | -0.5663410917 | 0.1139203722 | 33 | -4.971376769 | 2.01E-05 | 9.84E-05 |
| 1.94 | acetate - butyrate | 0.5296717917 | 0.1139203722 | 33 | 4.649491409 | 5.16E-05 | 2.21E-04 |
| 1.94 | propionate - butyrate | 1.096012883 | 0.1139203722 | 33 | 9.620868178 | 4.22E-11 | 5.79E-10 |
| 1.95 | acetate - propionate | -0.555568475 | 0.1168796187 | 33 | -4.753339216 | 3.81E-05 | 1.69E-04 |
| 1.95 | acetate - butyrate | 0.561004925 | 0.1168796187 | 33 | 4.799852458 | 3.32E-05 | 1.51E-04 |
| 1.95 | propionate - butyrate | 1.1165734 | 0.1168796187 | 33 | 9.553191673 | 5.02E-11 | 6.58E-10 |
| 1.96 | acetate - propionate | -0.5457207583 | 0.1186775241 | 33 | -4.59834971 | 5.99E-05 | 2.49E-04 |
| 1.96 | acetate - butyrate | 0.59622185 | 0.1186775241 | 33 | 5.023881773 | 1.72E-05 | 8.64E-05 |
| 1.96 | propionate - butyrate | 1.141942608 | 0.1186775241 | 33 | 9.622231483 | 4.21E-11 | 5.79E-10 |
| 1.97 | acetate - propionate | -0.54305585 | 0.1211841357 | 33 | -4.481245396 | 8.43E-05 | 3.34E-04 |
| 1.97 | acetate - butyrate | 0.6282967667 | 0.1211841357 | 33 | 5.18464536 | 1.07E-05 | 5.81E-05 |
| 1.97 | propionate - butyrate | 1.171352617 | 0.1211841357 | 33 | 9.665890756 | 3.77E-11 | 5.76E-10 |
| 1.98 | acetate - propionate | -0.54659545 | 0.1249113181 | 33 | -4.375868084 | 1.14E-04 | 4.08E-04 |

| | | | | | | | |
|---|---|---|---|---|---|---|---|
| 1.98 | acetate - butyrate | 0.6551515667 | 0.1249113181 | 33 | 5.244933581 | 8.96E-06 | 5.10E-05 |
| 1.98 | propionate - butyrate | 1.201747017 | 0.1249113181 | 33 | 9.620801665 | 4.22E-11 | 5.79E-10 |
| 1.99 | acetate - propionate | -0.5575298417 | 0.1285741891 | 33 | -4.336250109 | 1.28E-04 | 4.42E-04 |
| 1.99 | acetate - butyrate | 0.6872086167 | 0.1285741891 | 33 | 5.344841147 | 6.67E-06 | 3.96E-05 |
| 1.99 | propionate - butyrate | 1.244738458 | 0.1285741891 | 33 | 9.681091256 | 3.62E-11 | 5.75E-10 |
| 2 | acetate - propionate | -0.5728766917 | 0.130389777 | 33 | -4.393570607 | 1.09E-04 | 3.98E-04 |
| 2 | acetate - butyrate | 0.7135080417 | 0.130389777 | 33 | 5.472116435 | 4.58E-06 | 3.00E-05 |
| 2 | propionate - butyrate | 1.286384733 | 0.130389777 | 33 | 9.865687043 | 2.27E-11 | 4.17E-10 |
| 2.01 | acetate - propionate | -0.58905265 | 0.1358207091 | 33 | -4.336987001 | 1.28E-04 | 4.42E-04 |
| 2.01 | acetate - butyrate | 0.7482216917 | 0.1358207091 | 33 | 5.508892542 | 4.11E-06 | 2.78E-05 |
| 2.01 | propionate - butyrate | 1.337274342 | 0.1358207091 | 33 | 9.845879543 | 2.39E-11 | 4.23E-10 |
| 2.02 | acetate - propionate | -0.605746675 | 0.1403171576 | 33 | -4.316982222 | 1.36E-04 | 4.65E-04 |
| 2.02 | acetate - butyrate | 0.7894497 | 0.1403171576 | 33 | 5.626180813 | 2.90E-06 | 2.19E-05 |
| 2.02 | propionate - butyrate | 1.395196375 | 0.1403171576 | 33 | 9.943163035 | 1.87E-11 | 3.88E-10 |
| 2.03 | acetate - propionate | -0.6346480917 | 0.146063511 | 33 | -4.345014628 | 1.25E-04 | 4.38E-04 |
| 2.03 | acetate - butyrate | 0.8232612083 | 0.146063511 | 33 | 5.636323562 | 2.82E-06 | 2.15E-05 |
| 2.03 | propionate - butyrate | 1.4579093 | 0.146063511 | 33 | 9.98133819 | 1.70E-11 | 3.65E-10 |
| 2.04 | acetate - propionate | -0.6700786833 | 0.150999993 | 33 | -4.437607381 | 9.57E-05 | 3.61E-04 |
| 2.04 | acetate - butyrate | 0.8465632917 | 0.150999993 | 33 | 5.606379676 | 3.08E-06 | 2.29E-05 |
| 2.04 | propionate - butyrate | 1.516641975 | 0.150999993 | 33 | 10.04398706 | 1.45E-11 | 3.50E-10 |

| | | | | | | | |
|---|---|---|---|---|---|---|---|
| 2.05 | acetate - propionate | -0.718555125 | 0.1571704504 | 33 | -4.57182074 | 6.48E-05 | 2.66E-04 |
| 2.05 | acetate - butyrate | 0.8635089083 | 0.1571704504 | 33 | 5.494091961 | 4.29E-06 | 2.87E-05 |
| 2.05 | propionate - butyrate | 1.582064033 | 0.1571704504 | 33 | 10.0659127 | 1.37E-11 | 3.45E-10 |
| 2.06 | acetate - propionate | -0.7753290667 | 0.1654774556 | 33 | -4.68540602 | 4.65E-05 | 2.03E-04 |
| 2.06 | acetate - butyrate | 0.8789267917 | 0.1654774556 | 33 | 5.311459428 | 7.36E-06 | 4.31E-05 |
| 2.06 | propionate - butyrate | 1.654255858 | 0.1654774556 | 33 | 9.996865449 | 1.63E-11 | 3.64E-10 |
| 2.07 | acetate - propionate | -0.8392451083 | 0.1736702018 | 33 | -4.832407055 | 3.02E-05 | 1.40E-04 |
| 2.07 | acetate - butyrate | 0.8977650583 | 0.1736702018 | 33 | 5.169367278 | 1.12E-05 | 6.03E-05 |
| 2.07 | propionate - butyrate | 1.737010167 | 0.1736702018 | 33 | 10.00177433 | 1.61E-11 | 3.64E-10 |
| 2.08 | acetate - propionate | -0.9085375417 | 0.1848150684 | 33 | -4.915927848 | 2.36E-05 | 1.13E-04 |
| 2.08 | acetate - butyrate | 0.9143399667 | 0.1848150684 | 33 | 4.947323692 | 2.15E-05 | 1.04E-04 |
| 2.08 | propionate - butyrate | 1.822877508 | 0.1848150684 | 33 | 9.86325154 | 2.28E-11 | 4.17E-10 |
| 2.09 | acetate - propionate | -0.9914065667 | 0.1960000584 | 33 | -5.058195262 | 1.55E-05 | 7.98E-05 |
| 2.09 | acetate - butyrate | 0.915651675 | 0.1960000584 | 33 | 4.671690828 | 4.84E-05 | 2.08E-04 |
| 2.09 | propionate - butyrate | 1.907058242 | 0.1960000584 | 33 | 9.72988609 | 3.20E-11 | 5.21E-10 |
| 2.1 | acetate - propionate | -1.088450267 | 0.2089900934 | 33 | -5.208142878 | 9.98E-06 | 5.52E-05 |
| 2.1 | acetate - butyrate | 0.913805975 | 0.2089900934 | 33 | 4.372484647 | 1.16E-04 | 4.08E-04 |
| 2.1 | propionate - butyrate | 2.002256242 | 0.2089900934 | 33 | 9.580627525 | 4.68E-11 | 6.27E-10 |
| 2.11 | acetate - propionate | -1.196111525 | 0.2238581993 | 33 | -5.343166025 | 6.70E-06 | 3.96E-05 |
| 2.11 | acetate - butyrate | 0.9005234167 | 0.2238581993 | 33 | 4.022740375 | 3.15E-04 | 9.91E-04 |

| | | | | | | | |
|---|---|---|---|---|---|---|---|
| 2.11 | propionate - butyrate | 2.096634942 | 0.2238581993 | 33 | 9.3659064 | 8.13E-11 | 1.04E-09 |
| 2.12 | acetate - propionate | -1.303371225 | 0.2409131692 | 33 | -5.410128592 | 5.50E-06 | 3.53E-05 |
| 2.12 | acetate - butyrate | 0.8903455417 | 0.2409131692 | 33 | 3.695711383 | 7.91E-04 | 0.002292854687 |
| 2.12 | propionate - butyrate | 2.193716767 | 0.2409131692 | 33 | 9.105839975 | 1.60E-10 | 2.01E-09 |
| 2.13 | acetate - propionate | -1.422876925 | 0.2580320038 | 33 | -5.51434281 | 4.04E-06 | 2.78E-05 |
| 2.13 | acetate - butyrate | 0.8727985917 | 0.2580320038 | 33 | 3.38252069 | 0.00186392067 | 0.005040108359 |
| 2.13 | propionate - butyrate | 2.295675517 | 0.2580320038 | 33 | 8.8968635 | 2.78E-10 | 3.42E-09 |
| 2.14 | acetate - propionate | -1.54897815 | 0.2776538083 | 33 | -5.578811108 | 3.34E-06 | 2.43E-05 |
| 2.14 | acetate - butyrate | 0.8580636917 | 0.2776538083 | 33 | 3.090408508 | 0.004042240095 | 0.01024147386 |
| 2.14 | propionate - butyrate | 2.407041842 | 0.2776538083 | 33 | 8.669219616 | 5.09E-10 | 6.02E-09 |
| 2.15 | acetate - propionate | -1.678098058 | 0.3017076791 | 33 | -5.561999824 | 3.51E-06 | 2.52E-05 |
| 2.15 | acetate - butyrate | 0.8396317083 | 0.3017076791 | 33 | 2.782931183 | 0.008843033985 | 0.02066802129 |
| 2.15 | propionate - butyrate | 2.517729767 | 0.3017076791 | 33 | 8.344931007 | 1.22E-09 | 1.36E-08 |
| 2.16 | acetate - propionate | -1.817228117 | 0.3285834977 | 33 | -5.530491121 | 3.85E-06 | 2.73E-05 |
| 2.16 | acetate - butyrate | 0.813903575 | 0.3285834977 | 33 | 2.477006851 | 0.01854524149 | 0.04051732108 |
| 2.16 | propionate - butyrate | 2.631131692 | 0.3285834977 | 33 | 8.007497972 | 3.08E-09 | 3.31E-08 |
| 2.17 | acetate - propionate | -1.95576225 | 0.3579579865 | 33 | -5.463664239 | 4.69E-06 | 3.04E-05 |
| 2.17 | acetate - butyrate | 0.7932202333 | 0.3579579865 | 33 | 2.215959032 | 0.03370254896 | 0.06842638727 |
| 2.17 | propionate - butyrate | 2.748982483 | 0.3579579865 | 33 | 7.67962327 | 7.66E-09 | 7.82E-08 |
| 2.18 | acetate - propionate | -2.106148325 | 0.3897665496 | 33 | -5.403614874 | 5.60E-06 | 3.56E-05 |

| | | | | | | | |
|---|---|---|---|---|---|---|---|
| 2.18 | acetate - butyrate | 0.7656171333 | 0.3897665496 | 33 | 1.964296664 | 0.05796381988 | 0.1106081753 |
| 2.18 | propionate - butyrate | 2.871765458 | 0.3897665496 | 33 | 7.367911538 | 1.84E-08 | 1.79E-07 |
| 2.19 | acetate - propionate | -2.254747067 | 0.4213203383 | 33 | -5.351621704 | 6.54E-06 | 3.94E-05 |
| 2.19 | acetate - butyrate | 0.7256173167 | 0.4213203383 | 33 | 1.722246117 | 0.09438673135 | 0.1709165135 |
| 2.19 | propionate - butyrate | 2.980364383 | 0.4213203383 | 33 | 7.073867821 | 4.25E-08 | 3.95E-07 |
| 2.2 | acetate - propionate | -2.397387658 | 0.4591040743 | 33 | -5.221882776 | 9.59E-06 | 5.35E-05 |
| 2.2 | acetate - butyrate | 0.7025858417 | 0.4591040743 | 33 | 1.530341116 | 0.1354630581 | 0.232718587 |
| 2.2 | propionate - butyrate | 3.0999735 | 0.4591040743 | 33 | 6.752223893 | 1.07E-07 | 9.81E-07 |
| 2.21 | acetate - propionate | -2.534192042 | 0.4990672786 | 33 | -5.077856534 | 1.47E-05 | 7.69E-05 |
| 2.21 | acetate - butyrate | 0.6941576417 | 0.4990672786 | 33 | 1.390909946 | 0.1735610194 | 0.2899094036 |
| 2.21 | propionate - butyrate | 3.228349683 | 0.4990672786 | 33 | 6.46876648 | 2.44E-07 | 2.08E-06 |
| 2.22 | acetate - propionate | -2.692415325 | 0.5428344891 | 33 | -4.959919421 | 2.08E-05 | 1.01E-04 |
| 2.22 | acetate - butyrate | 0.6642088583 | 0.5428344891 | 33 | 1.223593695 | 0.2297708002 | 0.3694714466 |
| 2.22 | propionate - butyrate | 3.356624183 | 0.5428344891 | 33 | 6.183513116 | 5.63E-07 | 4.65E-06 |
| 2.23 | acetate - propionate | -2.826623825 | 0.5898113628 | 33 | -4.792420091 | 3.40E-05 | 1.53E-04 |
| 2.23 | acetate - butyrate | 0.641524475 | 0.5898113628 | 33 | 1.087677375 | 0.2846180966 | 0.4378181435 |
| 2.23 | propionate - butyrate | 3.4681483 | 0.5898113628 | 33 | 5.880097467 | 1.37E-06 | 1.08E-05 |
| 2.24 | acetate - propionate | -2.946605492 | 0.6382857781 | 33 | -4.616436074 | 5.69E-05 | 2.38E-04 |
| 2.24 | acetate - butyrate | 0.6297943083 | 0.6382857781 | 33 | 0.9866964452 | 0.3309711362 | 0.4915243565 |
| 2.24 | propionate - butyrate | 3.5763998 | 0.6382857781 | 33 | 5.603132519 | 3.11E-06 | 2.29E-05 |

| | | | | | | | |
|---|---|---|---|---|---|---|---|
| 2.25 | acetate - propionate | -3.083396142 | 0.6897865943 | 33 | -4.47007258 | 8.71E-05 | 3.39E-04 |
| 2.25 | acetate - butyrate | 0.6082301167 | 0.6897865943 | 33 | 0.8817656383 | 0.3842812908 | 0.5375613699 |
| 2.25 | propionate - butyrate | 3.691626258 | 0.6897865943 | 33 | 5.351838219 | 6.53E-06 | 3.94E-05 |
| 2.26 | acetate - propionate | -3.170412183 | 0.7420655739 | 33 | -4.272415127 | 1.54E-04 | 5.14E-04 |
| 2.26 | acetate - butyrate | 0.6117690083 | 0.7420655739 | 33 | 0.8244136769 | 0.4156238461 | 0.560262077 |
| 2.26 | propionate - butyrate | 3.782181192 | 0.7420655739 | 33 | 5.096828804 | 1.39E-05 | 7.33E-05 |
| 2.27 | acetate - propionate | -3.256288075 | 0.7938023864 | 33 | -4.102139438 | 2.52E-04 | 8.03E-04 |
| 2.27 | acetate - butyrate | 0.60303565 | 0.7938023864 | 33 | 0.759679815 | 0.452839139 | 0.5936130453 |
| 2.27 | propionate - butyrate | 3.859323725 | 0.7938023864 | 33 | 4.861819253 | 2.77E-05 | 1.29E-04 |
| 2.28 | acetate - propionate | -3.32115225 | 0.8501110341 | 33 | -3.906727612 | 4.38E-04 | 0.00132750205 |
| 2.28 | acetate - butyrate | 0.6137574417 | 0.8501110341 | 33 | 0.7219732683 | 0.4753960362 | 0.6164813115 |
| 2.28 | propionate - butyrate | 3.934909692 | 0.8501110341 | 33 | 4.62870088 | 5.49E-05 | 2.31E-04 |
| 2.29 | acetate - propionate | -3.341957742 | 0.9068395523 | 33 | -3.685280084 | 8.14E-04 | 0.002348906512 |
| 2.29 | acetate - butyrate | 0.6836576417 | 0.9068395523 | 33 | 0.7538904098 | 0.4562608975 | 0.5968011305 |
| 2.29 | propionate - butyrate | 4.025615383 | 0.9068395523 | 33 | 4.439170494 | 9.53E-05 | 3.61E-04 |
| 2.3 | acetate - propionate | -3.334598633 | 0.9588433832 | 33 | -3.477730244 | 0.001440238701 | 0.004002137956 |
| 2.3 | acetate - butyrate | 0.73621515 | 0.9588433832 | 33 | 0.7678158529 | 0.4480561399 | 0.5899079746 |
| 2.3 | propionate - butyrate | 4.070813783 | 0.9588433832 | 33 | 4.245546097 | 1.67E-04 | 5.46E-04 |
| 2.31 | acetate - propionate | -3.343267858 | 1.016365658 | 33 | -3.289434105 | 0.002392345303 | 0.006338921411 |
| 2.31 | acetate - butyrate | 0.7406992333 | 1.016365658 | 33 | 0.7287723936 | 0.4712815651 | 0.612462896 |

| | | | | | | | |
|---|---|---|---|---|---|---|---|
| 2.31 | propionate - butyrate | 4.083967092 | 1.016365658 | 33 | 4.018206498 | 3.19E-04 | 9.98E-04 |
| 2.32 | acetate - propionate | -3.269777317 | 1.072382127 | 33 | -3.049078527 | 0.004499903225 | 0.01121256878 |
| 2.32 | acetate - butyrate | 0.8379007833 | 1.072382127 | 33 | 0.7813453453 | 0.4401692552 | 0.58463009 |
| 2.32 | propionate - butyrate | 4.1076781 | 1.072382127 | 33 | 3.830423873 | 5.43E-04 | 0.00162904436 |
| 2.33 | acetate - propionate | -3.1890176 | 1.127210831 | 33 | -2.829122567 | 0.007880145248 | 0.01870758892 |
| 2.33 | acetate - butyrate | 0.9078092917 | 1.127210831 | 33 | 0.8053589149 | 0.4263776752 | 0.5726185704 |
| 2.33 | propionate - butyrate | 4.096826892 | 1.127210831 | 33 | 3.634481482 | 9.37E-04 | 0.002665270455 |
| 2.34 | acetate - propionate | -3.041008925 | 1.180796842 | 33 | -2.575387074 | 0.01468134735 | 0.03291023216 |
| 2.34 | acetate - butyrate | 1.04074845 | 1.180796842 | 33 | 0.8813950146 | 0.3844788613 | 0.5375613699 |
| 2.34 | propionate - butyrate | 4.081757375 | 1.180796842 | 33 | 3.456782089 | 0.001524647093 | 0.004198000901 |
| 2.35 | acetate - propionate | -2.876446642 | 1.232377149 | 33 | -2.334063598 | 0.02582935361 | 0.05408020911 |
| 2.35 | acetate - butyrate | 1.176516033 | 1.232377149 | 33 | 0.9546720617 | 0.3466849762 | 0.5061768539 |
| 2.35 | propionate - butyrate | 4.052962675 | 1.232377149 | 33 | 3.28873566 | 0.002396806106 | 0.006338921411 |
| 2.36 | acetate - propionate | -2.695127208 | 1.283787213 | 33 | -2.099356638 | 0.04351505873 | 0.08631440925 |
| 2.36 | acetate - butyrate | 1.302232358 | 1.283787213 | 33 | 1.014367759 | 0.3177871936 | 0.4754979597 |
| 2.36 | propionate - butyrate | 3.997359567 | 1.283787213 | 33 | 3.113724397 | 0.003803909978 | 0.009678302603 |
| 2.37 | acetate - propionate | -2.421840325 | 1.331649312 | 33 | -1.818677262 | 0.07804648018 | 0.1447425188 |
| 2.37 | acetate - butyrate | 1.46144635 | 1.331649312 | 33 | 1.097470886 | 0.2803793381 | 0.4335095921 |
| 2.37 | propionate - butyrate | 3.883286675 | 1.331649312 | 33 | 2.916148147 | 0.006327174708 | 0.0152611454 |
| 2.38 | acetate - propionate | -2.131058833 | 1.370567188 | 33 | -1.55487367 | 0.129515515 | 0.2231367301 |

| | | | | | | | |
|---|---|---|---|---|---|---|---|
| 2.38 | acetate - butyrate | 1.606478325 | 1.370567188 | 33 | 1.172126649 | 0.2495389919 | 0.3949178306 |
| 2.38 | propionate - butyrate | 3.737537158 | 1.370567188 | 33 | 2.727000318 | 0.01015588845 | 0.02364479049 |
| 2.39 | acetate - propionate | -1.827398592 | 1.417329663 | 33 | -1.289325017 | 0.206249349 | 0.3343235415 |
| 2.39 | acetate - butyrate | 1.794123458 | 1.417329663 | 33 | 1.265847675 | 0.2144316618 | 0.3466549385 |
| 2.39 | propionate - butyrate | 3.62152205 | 1.417329663 | 33 | 2.555172691 | 0.0154089058 | 0.03428623689 |
| 2.4 | acetate - propionate | -1.493488875 | 1.462789543 | 33 | -1.020986841 | 0.3146876144 | 0.4720314217 |
| 2.4 | acetate - butyrate | 1.997308708 | 1.462789543 | 33 | 1.365410846 | 0.1813599237 | 0.3012673112 |
| 2.4 | propionate - butyrate | 3.490797583 | 1.462789543 | 33 | 2.386397687 | 0.02290533693 | 0.04880536455 |
| 2.41 | acetate - propionate | -1.068677575 | 1.500526333 | 33 | -0.7122018131 | 0.4813451478 | 0.6228564896 |
| 2.41 | acetate - butyrate | 2.194551892 | 1.500526333 | 33 | 1.462521412 | 0.1530577506 | 0.2578039766 |
| 2.41 | propionate - butyrate | 3.263229467 | 1.500526333 | 33 | 2.174723225 | 0.0369205666 | 0.07396379288 |
| 2.42 | acetate - propionate | -0.6711738833 | 1.530622641 | 33 | -0.4384972922 | 0.663883225 | 0.7911493768 |
| 2.42 | acetate - butyrate | 2.293122492 | 1.530622641 | 33 | 1.498163186 | 0.1435964627 | 0.2432265926 |
| 2.42 | propionate - butyrate | 2.964296375 | 1.530622641 | 33 | 1.936660478 | 0.06138843179 | 0.1167735785 |
| 2.43 | acetate - propionate | -0.2472182333 | 1.563971158 | 33 | -0.1580708391 | 0.8753643022 | 0.9283587409 |
| 2.43 | acetate - butyrate | 2.479358925 | 1.563971158 | 33 | 1.585297089 | 0.1224362984 | 0.2127639422 |
| 2.43 | propionate - butyrate | 2.726577158 | 1.563971158 | 33 | 1.743367929 | 0.09058086082 | 0.1645188526 |
| 2.44 | acetate - propionate | 0.1636943417 | 1.591773993 | 33 | 0.1028376782 | 0.9187142003 | 0.9502107697 |
| 2.44 | acetate - butyrate | 2.5701327 | 1.591773993 | 33 | 1.614634182 | 0.1159124893 | 0.2037761837 |
| 2.44 | propionate - butyrate | 2.406438358 | 1.591773993 | 33 | 1.511796504 | 0.1401038162 | 0.2386514157 |

| | | | | | | | |
|---|---|---|---|---|---|---|---|
| 2.45 | acetate - propionate | 0.6671473667 | 1.620245225 | 33 | 0.4117570331 | 0.6831798239 | 0.8031132307 |
| 2.45 | acetate - butyrate | 2.784871708 | 1.620245225 | 33 | 1.71879643 | 0.09502088307 | 0.1715496781 |
| 2.45 | propionate - butyrate | 2.117724342 | 1.620245225 | 33 | 1.307039397 | 0.2002341538 | 0.3272119099 |
| 2.46 | acetate - propionate | 1.0432922 | 1.63710633 | 33 | 0.637278215 | 0.5283405455 | 0.6693053549 |
| 2.46 | acetate - butyrate | 2.956837908 | 1.63710633 | 33 | 1.806136751 | 0.08002520007 | 0.1475694056 |
| 2.46 | propionate - butyrate | 1.913545708 | 1.63710633 | 33 | 1.168858536 | 0.25083504 | 0.3949178306 |
| 2.47 | acetate - propionate | 1.323534458 | 1.651332649 | 33 | 0.8014947558 | 0.428579003 | 0.5742958641 |
| 2.47 | acetate - butyrate | 3.079910925 | 1.651332649 | 33 | 1.865106299 | 0.07107945213 | 0.133108415 |
| 2.47 | propionate - butyrate | 1.756376467 | 1.651332649 | 33 | 1.063611543 | 0.295226084 | 0.450686908 |
| 2.48 | acetate - propionate | 1.639846175 | 1.664704163 | 33 | 0.9850676242 | 0.3317585623 | 0.4915243565 |
| 2.48 | acetate - butyrate | 3.182553442 | 1.664704163 | 33 | 1.911783194 | 0.06461964497 | 0.1217676435 |
| 2.48 | propionate - butyrate | 1.542707267 | 1.664704163 | 33 | 0.9267155695 | 0.3608026344 | 0.5242505748 |
| 2.49 | acetate - propionate | 1.843865333 | 1.660702155 | 33 | 1.110292612 | 0.2748980017 | 0.4261272365 |
| 2.49 | acetate - butyrate | 3.301248225 | 1.660702155 | 33 | 1.987862914 | 0.05517592058 | 0.1062973805 |
| 2.49 | propionate - butyrate | 1.457382892 | 1.660702155 | 33 | 0.8775703018 | 0.3865215175 | 0.5376249795 |
| 2.5 | acetate - propionate | 1.89973335 | 1.651477267 | 33 | 1.150323646 | 0.258278503 | 0.4055779618 |
| 2.5 | acetate - butyrate | 3.3919063 | 1.651477267 | 33 | 2.053861938 | 0.04798159153 | 0.09363397959 |
| 2.5 | propionate - butyrate | 1.49217295 | 1.651477267 | 33 | 0.9035382927 | 0.372789 | 0.5356130702 |
| 2.51 | acetate - propionate | 1.915017508 | 1.636587277 | 33 | 1.170128556 | 0.2503308004 | 0.3949178306 |
| 2.51 | acetate - butyrate | 3.5025838 | 1.636587277 | 33 | 2.140175381 | 0.03982439843 | 0.07951692799 |

| | | | | | | | |
|---|---|---|---|---|---|---|---|
| 2.51 | propionate - butyrate | 1.587566292 | 1.636587277 | 33 | 0.9700468246 | 0.3390797686 | 0.4969854478 |
| 2.52 | acetate - propionate | 1.828453992 | 1.620332813 | 33 | 1.128443476 | 0.2672699533 | 0.4164438807 |
| 2.52 | acetate - butyrate | 3.567035617 | 1.620332813 | 33 | 2.201421577 | 0.03480725732 | 0.07043213477 |
| 2.52 | propionate - butyrate | 1.738581625 | 1.620332813 | 33 | 1.072978101 | 0.2910649006 | 0.4465957635 |
| 2.53 | acetate - propionate | 1.747187783 | 1.597690482 | 33 | 1.093570878 | 0.2820619087 | 0.4349957312 |
| 2.53 | acetate - butyrate | 3.582377533 | 1.597690482 | 33 | 2.242222492 | 0.03178602096 | 0.06497278183 |
| 2.53 | propionate - butyrate | 1.83518975 | 1.597690482 | 33 | 1.148651614 | 0.2589577753 | 0.4055884117 |
| 2.54 | acetate - propionate | 1.64422035 | 1.571039923 | 33 | 1.046580883 | 0.3028984963 | 0.4612318013 |
| 2.54 | acetate - butyrate | 3.576547417 | 1.571039923 | 33 | 2.276547759 | 0.02942871851 | 0.06056490534 |
| 2.54 | propionate - butyrate | 1.932327067 | 1.571039923 | 33 | 1.229966876 | 0.2274062326 | 0.3666469472 |
| 2.55 | acetate - propionate | 1.5463839 | 1.542337914 | 33 | 1.002623281 | 0.3233382475 | 0.4826063446 |
| 2.55 | acetate - butyrate | 3.540552175 | 1.542337914 | 33 | 2.295574882 | 0.02819086823 | 0.05841612901 |
| 2.55 | propionate - butyrate | 1.994168275 | 1.542337914 | 33 | 1.292951601 | 0.2050068363 | 0.3332051813 |
| 2.56 | acetate - propionate | 1.441292408 | 1.518845264 | 33 | 0.948939594 | 0.3495494302 | 0.509126344 |
| 2.56 | acetate - butyrate | 3.503436892 | 1.518845264 | 33 | 2.306645038 | 0.02749245397 | 0.05716534395 |
| 2.56 | propionate - butyrate | 2.062144483 | 1.518845264 | 33 | 1.357705444 | 0.1837694009 | 0.3027676196 |
| 2.57 | acetate - propionate | 1.353625383 | 1.49901052 | 33 | 0.9030125973 | 0.37306383 | 0.5356130702 |
| 2.57 | acetate - butyrate | 3.525629525 | 1.49901052 | 33 | 2.351971168 | 0.0247928221 | 0.05245639202 |
| 2.57 | propionate - butyrate | 2.172004142 | 1.49901052 | 33 | 1.448958571 | 0.1567858534 | 0.2633478262 |
| 2.58 | acetate - propionate | 1.310426908 | 1.474221626 | 33 | 0.8888941019 | 0.380493929 | 0.5375613699 |

| | | | | | | | |
|---|---|---|---|---|---|---|---|
| 2.58 | acetate - butyrate | 3.558697283 | 1.474221626 | 33 | 2.413949992 | 0.02148982093 | 0.04611516732 |
| 2.58 | propionate - butyrate | 2.248270375 | 1.474221626 | 33 | 1.525055891 | 0.1367728511 | 0.2343012194 |
| 2.59 | acetate - propionate | 1.288468717 | 1.455241603 | 33 | 0.8853984889 | 0.3823481468 | 0.5375613699 |
| 2.59 | acetate - butyrate | 3.591060175 | 1.455241603 | 33 | 2.467672836 | 0.01895648986 | 0.04097047809 |
| 2.59 | propionate - butyrate | 2.302591458 | 1.455241603 | 33 | 1.582274347 | 0.123125212 | 0.2133462725 |
| 2.6 | acetate - propionate | 1.229704633 | 1.424825122 | 33 | 0.8630565351 | 0.3943357093 | 0.5430132857 |
| 2.6 | acetate - butyrate | 3.46740795 | 1.424825122 | 33 | 2.433567387 | 0.02053100827 | 0.04421499282 |
| 2.6 | propionate - butyrate | 2.237703317 | 1.424825122 | 33 | 1.570510852 | 0.1258364202 | 0.2174193736 |
| 2.61 | acetate - propionate | 1.207787333 | 1.400221449 | 33 | 0.8625687987 | 0.3946000314 | 0.5430132857 |
| 2.61 | acetate - butyrate | 3.462007983 | 1.400221449 | 33 | 2.472471755 | 0.01874402348 | 0.04065700057 |
| 2.61 | propionate - butyrate | 2.25422065 | 1.400221449 | 33 | 1.609902956 | 0.1169448564 | 0.2043992707 |
| 2.62 | acetate - propionate | 1.194174458 | 1.372603991 | 33 | 0.8700065467 | 0.3905814285 | 0.5401848656 |
| 2.62 | acetate - butyrate | 3.475137733 | 1.372603991 | 33 | 2.531784663 | 0.01629201752 | 0.0361179653 |
| 2.62 | propionate - butyrate | 2.280963275 | 1.372603991 | 33 | 1.661778116 | 0.1060295366 | 0.1886012111 |
| 2.63 | acetate - propionate | 1.160042125 | 1.346965286 | 33 | 0.8612264453 | 0.3953280803 | 0.5430132857 |
| 2.63 | acetate - butyrate | 3.447852633 | 1.346965286 | 33 | 2.559718999 | 0.01524243972 | 0.0340414487 |
| 2.63 | propionate - butyrate | 2.287810508 | 1.346965286 | 33 | 1.698492554 | 0.09882603652 | 0.1773574405 |
| 2.64 | acetate - propionate | 1.147843542 | 1.338302478 | 33 | 0.857686181 | 0.3972522742 | 0.5438109661 |
| 2.64 | acetate - butyrate | 3.499534217 | 1.338302478 | 33 | 2.61490528 | 0.01334965624 | 0.03003672655 |
| 2.64 | propionate - butyrate | 2.351690675 | 1.338302478 | 33 | 1.757219099 | 0.08815580807 | 0.1605980431 |

| | | | | | | | |
|---|---|---|---|---|---|---|---|
| 2.65 | acetate - propionate | 1.161623158 | 1.327401814 | 33 | 0.8751104195 | 0.3878389155 | 0.5376249795 |
| 2.65 | acetate - butyrate | 3.545016667 | 1.327401814 | 33 | 2.670643229 | 0.01166046868 | 0.02663357051 |
| 2.65 | propionate - butyrate | 2.383393508 | 1.327401814 | 33 | 1.795532809 | 0.08173146879 | 0.1493456839 |
| 2.66 | acetate - propionate | 1.158022775 | 1.297690113 | 33 | 0.8923723491 | 0.3786546677 | 0.5375613699 |
| 2.66 | acetate - butyrate | 3.494265767 | 1.297690113 | 33 | 2.692681196 | 0.01104903092 | 0.02542963986 |
| 2.66 | propionate - butyrate | 2.336242992 | 1.297690113 | 33 | 1.800308847 | 0.08095917781 | 0.148455791 |
| 2.67 | acetate - propionate | 1.133182708 | 1.269751065 | 33 | 0.8924447785 | 0.3786164288 | 0.5375613699 |
| 2.67 | acetate - butyrate | 3.448829367 | 1.269751065 | 33 | 2.71614607 | 0.01043083107 | 0.02409881662 |
| 2.67 | propionate - butyrate | 2.315646658 | 1.269751065 | 33 | 1.823701292 | 0.07726552515 | 0.1437997274 |
| 2.68 | acetate - propionate | 1.125305867 | 1.255469551 | 33 | 0.8963227072 | 0.376572714 | 0.5375613699 |
| 2.68 | acetate - butyrate | 3.516679808 | 1.255469551 | 33 | 2.801087295 | 0.008452135068 | 0.01990874002 |
| 2.68 | propionate - butyrate | 2.391373942 | 1.255469551 | 33 | 1.904764588 | 0.06555740864 | 0.1231498985 |
| 2.69 | acetate - propionate | 1.113292608 | 1.228224716 | 33 | 0.9064242024 | 0.3712826041 | 0.5356062446 |
| 2.69 | acetate - butyrate | 3.528813225 | 1.228224716 | 33 | 2.873100647 | 0.007055416327 | 0.01688260335 |
| 2.69 | propionate - butyrate | 2.415520617 | 1.228224716 | 33 | 1.966676444 | 0.05767683733 | 0.1104099458 |
| 2.7 | acetate - propionate | 1.0887046 | 1.200525333 | 33 | 0.9068568321 | 0.3710571196 | 0.5356062446 |
| 2.7 | acetate - butyrate | 3.5269792 | 1.200525333 | 33 | 2.937863204 | 0.005987235483 | 0.01455767337 |
| 2.7 | propionate - butyrate | 2.4382746 | 1.200525333 | 33 | 2.031006372 | 0.05037395318 | 0.09735735181 |
| 2.71 | acetate - propionate | 1.071487925 | 1.193371518 | 33 | 0.8978661788 | 0.3757612702 | 0.5375613699 |
| 2.71 | acetate - butyrate | 3.60015975 | 1.193371518 | 33 | 3.016797112 | 0.004891066197 | 0.01208734802 |

| | | | | | | | |
|---|---|---|---|---|---|---|---|
| 2.71 | propionate - butyrate | 2.528671825 | 1.193371518 | 33 | 2.118930933 | 0.04170908516 | 0.08300520909 |
| 2.72 | acetate - propionate | 1.0377058 | 1.174782046 | 33 | 0.8833177211 | 0.3834546189 | 0.5375613699 |
| 2.72 | acetate - butyrate | 3.601189083 | 1.174782046 | 33 | 3.065410383 | 0.004313475906 | 0.01088295385 |
| 2.72 | propionate - butyrate | 2.563483283 | 1.174782046 | 33 | 2.182092662 | 0.03632610426 | 0.07325966845 |
| 2.73 | acetate - propionate | 1.010293067 | 1.147804952 | 33 | 0.8801957729 | 0.3851185933 | 0.5375613699 |
| 2.73 | acetate - butyrate | 3.60965825 | 1.147804952 | 33 | 3.144835927 | 0.003506621866 | 0.00895971604 |
| 2.73 | propionate - butyrate | 2.599365183 | 1.147804952 | 33 | 2.264640154 | 0.03022805638 | 0.06199836053 |
| 2.74 | acetate - propionate | 0.9591203833 | 1.12494369 | 33 | 0.8525941274 | 0.4000302322 | 0.5445106773 |
| 2.74 | acetate - butyrate | 3.6384554 | 1.12494369 | 33 | 3.234344469 | 0.002769711199 | 0.007261460231 |
| 2.74 | propionate - butyrate | 2.679335017 | 1.12494369 | 33 | 2.381750341 | 0.02315228009 | 0.04915783412 |
| 2.75 | acetate - propionate | 0.913937 | 1.107500865 | 33 | 0.825224638 | 0.4151699145 | 0.560262077 |
| 2.75 | acetate - butyrate | 3.680749175 | 1.107500865 | 33 | 3.323472959 | 0.002184323811 | 0.005853987813 |
| 2.75 | propionate - butyrate | 2.766812175 | 1.107500865 | 33 | 2.498248321 | 0.01763961802 | 0.03867887152 |
| 2.76 | acetate - propionate | 0.8675575583 | 1.084697822 | 33 | 0.7998149722 | 0.4295380921 | 0.5743048105 |
| 2.76 | acetate - butyrate | 3.727886058 | 1.084697822 | 33 | 3.43679685 | 0.00160960156 | 0.004391808781 |
| 2.76 | propionate - butyrate | 2.8603285 | 1.084697822 | 33 | 2.636981878 | 0.01265521168 | 0.02858087132 |
| 2.77 | acetate - propionate | 0.8293518083 | 1.069616918 | 33 | 0.7753727475 | 0.44364061 | 0.5879456876 |
| 2.77 | acetate - butyrate | 3.8222123 | 1.069616918 | 33 | 3.573440394 | 0.001108606767 | 0.003138450144 |
| 2.77 | propionate - butyrate | 2.992860492 | 1.069616918 | 33 | 2.798067646 | 0.008516007719 | 0.01998113873 |
| 2.78 | acetate - propionate | 0.79076765 | 1.060820548 | 33 | 0.7454301783 | 0.4612884668 | 0.6020713106 |

| | | | | | | | |
|---|---|---|---|---|---|---|---|
| 2.78 | acetate - butyrate | 3.903445525 | 1.060820548 | 33 | 3.679647358 | 8.27E-04 | 0.002374525844 |
| 2.78 | propionate - butyrate | 3.112677875 | 1.060820548 | 33 | 2.934217179 | 0.006043084316 | 0.0146344572 |
| 2.79 | acetate - propionate | 0.7022086167 | 1.044768719 | 33 | 0.6721187223 | 0.5061869601 | 0.6483354266 |
| 2.79 | acetate - butyrate | 3.897130608 | 1.044768719 | 33 | 3.730137146 | 7.19E-04 | 0.002103800201 |
| 2.79 | propionate - butyrate | 3.194921992 | 1.044768719 | 33 | 3.058018423 | 0.004396929519 | 0.01104728542 |
| 2.8 | acetate - propionate | 0.6191396417 | 1.031789255 | 33 | 0.6000640523 | 0.5525622079 | 0.6927131214 |
| 2.8 | acetate - butyrate | 3.91708165 | 1.031789255 | 33 | 3.796397016 | 5.97E-04 | 0.001774374219 |
| 2.8 | propionate - butyrate | 3.297942008 | 1.031789255 | 33 | 3.196332963 | 0.003062543348 | 0.00795997258 |
| 2.81 | acetate - propionate | 0.5583937083 | 1.01786391 | 33 | 0.5485936804 | 0.5869770156 | 0.7312957446 |
| 2.81 | acetate - butyrate | 3.990684608 | 1.01786391 | 33 | 3.920646533 | 4.21E-04 | 0.001282804648 |
| 2.81 | propionate - butyrate | 3.4322909 | 1.01786391 | 33 | 3.372052853 | 0.001917221975 | 0.005161093084 |
| 2.82 | acetate - propionate | 0.4693105083 | 1.012889506 | 33 | 0.4633383064 | 0.6461633509 | 0.7792730012 |
| 2.82 | acetate - butyrate | 4.0233389 | 1.012889506 | 33 | 3.972139978 | 3.64E-04 | 0.001120229242 |
| 2.82 | propionate - butyrate | 3.554028392 | 1.012889506 | 33 | 3.508801671 | 0.001323279344 | 0.003711336952 |
| 2.83 | acetate - propionate | 0.3970963083 | 1.00217846 | 33 | 0.3962331304 | 0.6944835775 | 0.8100069578 |
| 2.83 | acetate - butyrate | 4.071165867 | 1.00217846 | 33 | 4.062316274 | 2.82E-04 | 8.94E-04 |
| 2.83 | propionate - butyrate | 3.674069558 | 1.00217846 | 33 | 3.666083144 | 8.59E-04 | 0.002453790657 |
| 2.84 | acetate - propionate | 0.3197204833 | 0.9925736244 | 33 | 0.3221126126 | 0.7493991422 | 0.8473517595 |
| 2.84 | acetate - butyrate | 4.13101995 | 0.9925736244 | 33 | 4.161927991 | 2.12E-04 | 6.87E-04 |
| 2.84 | propionate - butyrate | 3.811299467 | 0.9925736244 | 33 | 3.839815378 | 5.29E-04 | 0.001594595544 |

| | | | | | | | |
|---|---|---|---|---|---|---|---|
| 2.85 | acetate - propionate | 0.23993235 | 0.9894373929 | 33 | 0.2424937158 | 0.809898458 | 0.8911838872 |
| 2.85 | acetate - butyrate | 4.206107742 | 0.9894373929 | 33 | 4.251009484 | 1.64E-04 | 5.41E-04 |
| 2.85 | propionate - butyrate | 3.966175392 | 0.9894373929 | 33 | 4.008515769 | 3.28E-04 | 0.001015570899 |
| 2.86 | acetate - propionate | 0.1443453167 | 0.9858080356 | 33 | 0.1464233517 | 0.884478034 | 0.9356081308 |
| 2.86 | acetate - butyrate | 4.218066942 | 0.9858080356 | 33 | 4.278791397 | 1.52E-04 | 5.08E-04 |
| 2.86 | propionate - butyrate | 4.073721625 | 0.9858080356 | 33 | 4.132368045 | 2.31E-04 | 7.40E-04 |
| 2.87 | acetate - propionate | 0.05144936667 | 0.9845426851 | 33 | 0.05225712145 | 0.9586388763 | 0.9733061738 |
| 2.87 | acetate - butyrate | 4.248481358 | 0.9845426851 | 33 | 4.315182493 | 1.36E-04 | 4.65E-04 |
| 2.87 | propionate - butyrate | 4.197031992 | 0.9845426851 | 33 | 4.262925372 | 1.59E-04 | 5.25E-04 |
| 2.88 | acetate - propionate | -0.01851505833 | 0.9801500496 | 33 | -0.01889002438 | 0.9850425912 | 0.9899678042 |
| 2.88 | acetate - butyrate | 4.2929618 | 0.9801500496 | 33 | 4.37990265 | 1.13E-04 | 4.07E-04 |
| 2.88 | propionate - butyrate | 4.311476858 | 0.9801500496 | 33 | 4.398792675 | 1.07E-04 | 3.96E-04 |
| 2.89 | acetate - propionate | -0.119255775 | 0.9794506651 | 33 | -0.1217578172 | 0.9038292511 | 0.9429222118 |
| 2.89 | acetate - butyrate | 4.302914483 | 0.9794506651 | 33 | 4.39319165 | 1.09E-04 | 3.98E-04 |
| 2.89 | propionate - butyrate | 4.422170258 | 0.9794506651 | 33 | 4.514949467 | 7.64E-05 | 3.09E-04 |
| 2.9 | acetate - propionate | -0.22716335 | 0.9782389839 | 33 | -0.2322166196 | 0.8178038888 | 0.8949832031 |
| 2.9 | acetate - butyrate | 4.312855992 | 0.9782389839 | 33 | 4.40879587 | 1.04E-04 | 3.87E-04 |
| 2.9 | propionate - butyrate | 4.540019342 | 0.9782389839 | 33 | 4.641012489 | 5.29E-05 | 2.25E-04 |
| 2.91 | acetate - propionate | -0.3145130917 | 0.9786775375 | 33 | -0.3213653932 | 0.7499601584 | 0.8473517595 |
| 2.91 | acetate - butyrate | 4.3633807 | 0.9786775375 | 33 | 4.458445742 | 9.01E-05 | 3.48E-04 |

| | | | | | | | |
|---|---|---|---|---|---|---|---|
| 2.91 | propionate - butyrate | 4.677893792 | 0.9786775375 | 33 | 4.779811136 | 3.52E-05 | 1.57E-04 |
| 2.92 | acetate - propionate | -0.3980369667 | 0.9817248285 | 33 | -0.4054465723 | 0.6877660196 | 0.8058390972 |
| 2.92 | acetate - butyrate | 4.401793067 | 0.9817248285 | 33 | 4.483734076 | 8.37E-05 | 3.34E-04 |
| 2.92 | propionate - butyrate | 4.799830033 | 0.9817248285 | 33 | 4.889180648 | 2.56E-05 | 1.20E-04 |
| 2.93 | acetate - propionate | -0.5020939333 | 0.9823064585 | 33 | -0.5111377707 | 0.6126576197 | 0.7554857765 |
| 2.93 | acetate - butyrate | 4.397142575 | 0.9823064585 | 33 | 4.47634497 | 8.55E-05 | 3.36E-04 |
| 2.93 | propionate - butyrate | 4.899236508 | 0.9823064585 | 33 | 4.987482741 | 1.91E-05 | 9.46E-05 |
| 2.94 | acetate - propionate | -0.5978714667 | 0.9900050694 | 33 | -0.60390748 | 0.5500344525 | 0.690980781 |
| 2.94 | acetate - butyrate | 4.408291267 | 0.9900050694 | 33 | 4.45279666 | 9.16E-05 | 3.52E-04 |
| 2.94 | propionate - butyrate | 5.006162733 | 0.9900050694 | 33 | 5.05670414 | 1.56E-05 | 7.98E-05 |
| 2.95 | acetate - propionate | -0.6796198667 | 0.9930280031 | 33 | -0.6843914417 | 0.4985066403 | 0.6423066327 |
| 2.95 | acetate - butyrate | 4.416125383 | 0.9930280031 | 33 | 4.447130765 | 9.31E-05 | 3.55E-04 |
| 2.95 | propionate - butyrate | 5.09574525 | 0.9930280031 | 33 | 5.131522207 | 1.25E-05 | 6.68E-05 |
| 2.96 | acetate - propionate | -0.77051475 | 0.998264204 | 33 | -0.771854532 | 0.4456930785 | 0.5887030974 |
| 2.96 | acetate - butyrate | 4.417533842 | 0.998264204 | 33 | 4.425215112 | 9.92E-05 | 3.72E-04 |
| 2.96 | propionate - butyrate | 5.188048592 | 0.998264204 | 33 | 5.197069644 | 1.03E-05 | 5.66E-05 |
| 2.97 | acetate - propionate | -0.86045385 | 1.004216435 | 33 | -0.8568410351 | 0.3977124976 | 0.5438109661 |
| 2.97 | acetate - butyrate | 4.402565975 | 1.004216435 | 33 | 4.384080781 | 1.12E-04 | 4.06E-04 |
| 2.97 | propionate - butyrate | 5.263019825 | 1.004216435 | 33 | 5.240921816 | 9.06E-06 | 5.11E-05 |
| 2.98 | acetate - propionate | -0.9340799583 | 1.010915419 | 33 | -0.9239941743 | 0.3621967912 | 0.5250112142 |

| | | | | | | | |
|---|---|---|---|---|---|---|---|
| 2.98 | acetate - butyrate | 4.41946415 | 1.010915419 | 33 | 4.371744722 | 1.16E-04 | 4.08E-04 |
| 2.98 | propionate - butyrate | 5.353544108 | 1.010915419 | 33 | 5.295738897 | 7.71E-06 | 4.47E-05 |
| 2.99 | acetate - propionate | -0.9925813167 | 1.018955077 | 33 | -0.9741168562 | 0.3370853847 | 0.4969742958 |
| 2.99 | acetate - butyrate | 4.461769075 | 1.018955077 | 33 | 4.378769166 | 1.13E-04 | 4.07E-04 |
| 2.99 | propionate - butyrate | 5.454350392 | 1.018955077 | 33 | 5.352886022 | 6.51E-06 | 3.94E-05 |
| 3 | acetate - propionate | -1.07204325 | 1.0262563 | 33 | -1.044615512 | 0.3037927559 | 0.4614282917 |
| 3 | acetate - butyrate | 4.457769692 | 1.0262563 | 33 | 4.343719685 | 1.26E-04 | 4.38E-04 |
| 3 | propionate - butyrate | 5.529812942 | 1.0262563 | 33 | 5.388335197 | 5.86E-06 | 3.65E-05 |
